# Climbing fiber synapses rapidly inhibit neighboring Purkinje cells via ephaptic coupling

**DOI:** 10.1101/2019.12.17.879890

**Authors:** Kyung-Seok Han, Christopher H. Chen, Mehak M. Khan, Chong Guo, Wade G. Regehr

## Abstract

Climbing fibers (CFs) from the inferior olive (IO) provide strong excitatory inputs onto the dendrites of cerebellar Purkinje cells (PC), and trigger distinctive responses known as complex spikes (CSs). We find that in awake, behaving mice, a CS in one PC suppresses conventional simple spikes (SSs) in neighboring PCs for several milliseconds. This involves a novel form of ephaptic coupling, in which an excitatory synapse nonsynaptically inhibits neighboring cells by generating large negative extracellular signals near their dendrites. The distance dependence of CS-SS ephaptic signaling, combined with the known divergence of CF synapses made by IO neurons, allows a single IO neuron to influence the output of the cerebellum by synchronously suppressing the firing of potentially over one hundred PCs. Optogenetic studies *in vivo* and dynamic clamp studies in slice indicate that such brief PC suppression can effectively promote firing in neurons in the deep cerebellar nuclei and motor thalamus.

## Introduction

Climbing fiber (CF) synapses onto Purkinje cells (PCs) make essential contributions to cerebellar learning and cerebellar function. PCs are the sole outputs of the cerebellar cortex. Each PC receives a single powerful CF synapse from the inferior olive (Eccles et al., 1966; Szentágothai and Rajkovits, 1959). Activation of a CF onto a PC evokes a characteristic response known as a complex spike (CS) (Eccles et al., 1966; Fujita, 1968; Llinas and Sugimori, 1980) that arises from strong depolarization, dendritic calcium electrogenesis, and a sodium action potential followed by a series of spikelets (Davie et al., 2008; Eccles et al., 1966; Llinas and Nicholson, 1971; Llinas et al., 1968). CSs occur at 1-2 Hz and are readily distinguished from conventional sodium spikes known as simple spikes (SSs) (Eccles et al., 1966) that occur at frequencies of tens to over 100 Hz. Mossy fiber inputs to the cerebellar cortex activate granule cells (grCs), and tens of thousands of grCs form weak synapses onto each PC. It is thought that the cerebellum transforms mossy fiber inputs into PC outputs that serve as predictions of behavioral outcome (Medina, 2011). CFs provide instructive signals that regulate those grC to PC synapses that contribute to flawed predictions (Gilbert and Thach, 1977; Kitazawa et al., 1998; Lang et al., 2017). CSs are followed by a pause in SSs (Barmack and Yakhnitsa, 2003; Bell and Grimm, 1969; Granit and Phillips, 1956; Jin et al., 2017; Tang et al., 2017; Thach, 1967), allowing CFs to directly affect cerebellar output (Apps et al., 2018; Bengtsson et al., 2011; Blenkinsop and Lang, 2011; Eccles et al., 1966; Lang and Blenkinsop, 2011; Sudhakar et al., 2015; Tang et al., 2019; Tang et al., 2016; Yarden-Rabinowitz and Yarom, 2017). In addition, CF to PC synapses release so much glutamate that it slowly spills over to activate nearby molecular layer interneurons (MLIs) and Golgi cells (GCs) (Coddington et al., 2013; Eccles et al., 1966; Jorntell and Ekerot, 2003; Mathews et al., 2012; Nietz et al., 2017; Szapiro and Barbour, 2007). Thus, the CF plays a central role in cerebellar learning, regulates activity within the cerebellar cortex, and controls the output of the cerebellar cortex.

Here we ask whether CF synapses can directly inhibit nearby PCs via ephaptic signaling. Ephaptic signaling occurs when extracellular potentials directly influence firing (Anastassiou et al., 2015; Anastassiou et al., 2011; Blot and Barbour, 2014; Furukawa and Furshpan, 1963; Han et al., 2018; Korn and Axelrad, 1980; Korn and Faber, 1975; Weiss and Faber, 2010). Most ephaptic signaling described previously involves changes in extracellular potential near the axon initial segment where action potential firing is initiated. For example, at synapses between cerebellar basket cells and PCs, current flow through potassium channels of a presynaptic specialization known as a pinceau produces depolarizing extracellular signals near the axon that transiently inhibit the target PC (Blot and Barbour, 2014; Korn and Axelrad, 1980). Ephaptic coupling also occurs between PCs to promote synchronous firing. During a SS, sodium channels in the initial segment generate a hyperpolarizing extracellular signal that can directly open sodium channels in the axons of neighboring PCs to promote spike initiation (Han et al., 2018). We hypothesized that CSs could also generate large extracellular signals that would influence the firing of nearby PCs via ephaptic signaling.

We tested the hypothesis that CSs in one PC influence SS firing in neighboring PCs. Using *in vivo* multielectrode recordings, we found that CSs immediately suppress SSs in neighboring cells for about two milliseconds, after which SS firing in neighboring cells returns to baseline levels. We found that this suppression was a consequence of the complex spatiotemporal extracellular signals produced by CF activation. For CSs, the extracellular signals near the axons were not very effective at promoting SS firing in nearby PCs. Instead, the influence of the CF on nearby PCs was mediated by a novel form of ephaptic signaling whereby current flow through dendritic ionotropic glutamate receptors at a single CF synapse generates dendritically-localized extracellular hyperpolarization that reduces the firing of neighboring PCs. Based on the observed spread of ephaptic signaling (approximately 50 μm), the close spacing of PCs (Altman and Winfree, 1977), and the fact that IO neurons form CF synapses with multiple PCs (∼7 PCs) (Sugihara et al., 2001), our findings suggest that a single IO cell can synchronously suppress PC firing in over 100 PCs. Dynamic clamp and optogenetic studies indicate that such brief inhibition of PC firing leads to transient disinhibition that can be highly effective at promoting firing in target cells in the deep cerebellar nuclei.

## Results

In order to assess whether CSs in one cell influence SSs in neighboring cells, we recorded from pairs of Purkinje cells in awake, head-restrained animals on a cylindrical treadmill using a silicon probe with a linear array of electrodes (Figure 1A). Analysis was restricted to cases where the firing of individual PCs could be isolated on two or more electrodes. SSs occurred at much higher frequencies and had very different waveforms than CSs (Figure 1B). Following a CS there was a characteristic pause of SS firing in that same cell (Figure S2). To our surprise, CSs in one cell produced a short-latency transient decrease in SSs in a PC recorded on an adjacent site (Figures 1C-1G, S2B). CSs recorded on one electrode (Figure 1Ca) were used to align recordings from a second site (Figure 1Cb). It was immediately apparent from superimposed trials, (Figure 1Cb), a raster plot (Figure 1Cc), and a histogram (Figure 1Cd) that there were fewer spikes in PC2 immediately after the occurrence of the CS in PC1. This pause began at the onset of the CS in PC1, and persisted for approximately 2 ms (Figure 1Cd). Analysis of many pairs of PCs revealed that this decrease in firing was dependent on distance (Figure 1D). Suppression was largest for recordings from neighboring contacts (25 μm separation), was still apparent for cells separated by 50 μm, and absent for electrodes separated by 75 μm or more (Figure 1E).

**Figure 1.**
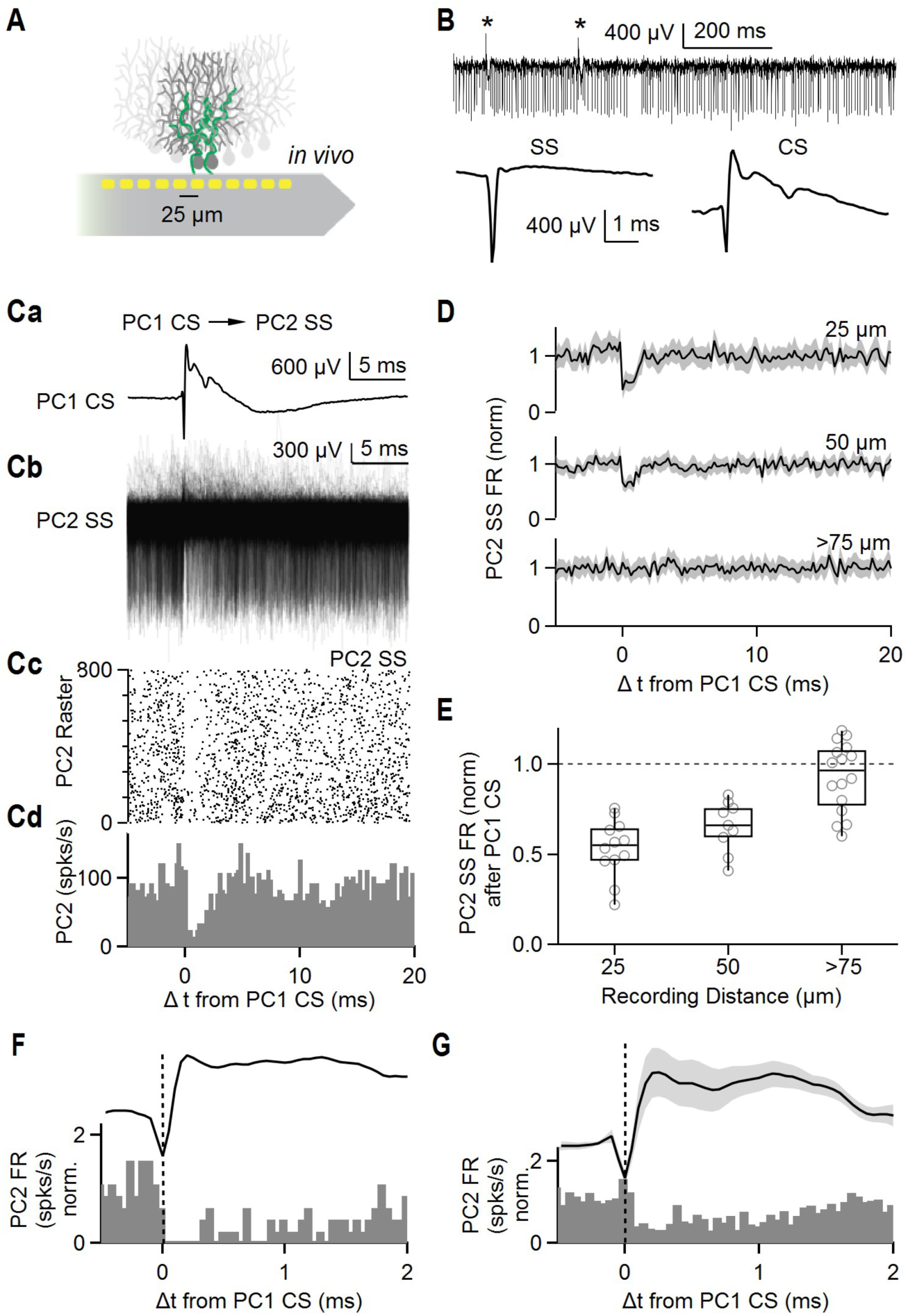
Complex spikes in a Purkinje cell rapidly and transiently suppress simple-spike firing in neighboring PCs in awake mice. (A) Schematic showing recording probe near PCs innervated by climbing fibers (green). (B) (top) Raw traces showing spikes recorded from single recording site. Asterisks indicate complex spikes. (bottom) Average simple spike (SS) and complex spike (CS), 10 minute recording. (C, a) Average complex spike recorded on single site (PC1 CS). (C, b) Simple spikes simultaneously recorded on a neighboring site (PC2 SS) are aligned to PC1 CS. (C, c) Raster plot of simple spikes from (C, b). (C, d) Histogram summarizing the data in (C, b) (D) Average firing rate of simple spikes from neighboring PCs (PC2 SS) after complex spikes from PC1 (PC1 CS). Recording sites were separated by 25 μm (top, n=12 pairs), 50 μm (middle, n=9), and more than 75 μm (bottom, n=16). Shaded grey is SEM. (E) Summary of normalized firing rates of PC2 SS after PC1 CS as a function of distance between recording sites. (F) The CS waveform and the histogram of SSs recorded in a neighboring cell from (C) shown on an expanded time scale. (G) The average CS waveform and the histogram of SSs recorded in nearest neighbor PCs are displayed to illustrate the relative timing.

A CS in one cell suppressed SSs in neighboring cells with remarkable speed. A comparison of the CS waveform and SS histogram for a neighboring PC illustrates that SS suppression in neighboring cells begins at approximately the same time as the peak of the negative component of the CS, well before the positive component of the CS (Figure 1F). The average CS waveform and the average SS responses of neighboring cells showed similar timing (Figure 1G). This observation constrains the mechanism that allows CSs to influence the firing of neighboring PCs.

The CS suppression of SSs in neighboring PCs raises two important questions. First, what is the mechanism? Second, what are the functional implications of this suppression? Based on the high packing density of PCs in the cerebellum (Altman and Winfree, 1977), and the fact that each IO neuron makes CF synapses with an average of 7 PCs, it is expected that a when a single IO neuron fires an action potential it could simultaneously suppress firing of up to 100 PCs. It is not known if transient PC suppression provides an effective means of activating targets in the deep cerebellar nuclei (DCN). We will begin by addressing the question of mechanism, and then return to the issue of functional significance.

### Mechanism of CS suppression of spiking in nearby PCs

We considered a number of possible mechanisms that could allow CSs to suppress SSs in neighboring cells. Gap junction coupling can allow neurons to inhibit the firing of neighboring cells, but this is not a viable mechanism because neighboring PCs are not electrically coupled (Han et al., 2018). Another possibility is that CF activation could evoke disynaptic inhibition in neighboring PCs. Both PCs and MLIs inhibit PCs (Witter et al., 2016); (Orduz and Llano, 2007); (Dizon and Khodakhah, 2011). However, although PCs make collateral synapses onto other PCs, it is very rare for a PC to contact its nearest neighbors (fewer than 10%, (Witter et al., 2016)). This connectivity is incompatible with our observation that PCs consistently suppress the firing of neighboring PCs. We also considered the possibility that CFs excite MLIs to produce rapid disynaptic inhibition. It is known that glutamate spillover from CF to PC synapses excites MLIs that in turn inhibit nearby PCs. However, the time course of CF→MLI→PC inhibition is not consistent with a short latency inhibition of PCs. The spillover current in MLIs begins 1-2 milliseconds after CF-EPSCs in PCs (Szapiro and Barbour, 2007, Figure 2), has a rise time of 0.7 ms (Coddington et al., 2013), and evokes IPSCs that begin approximately 3-5 milliseconds after CF-EPSC onset (Coddington et al., 2013, Figure 1G and text). This makes it unlikely that MLIs are involved in the rapid CS-evoked inhibition of nearby PCs. We also experimentally tested this possibility by examining the effect of MLI suppression on CS suppression of SSs on neighboring cells (Figure S3). We used conditional viruses in c-kit Cre mice to express halorhodopsin in MLIs but not PCs (Amat et al., 2017). We found that optical suppression of MLI firing elevated PC firing but did not prevent CSs from inhibiting neighboring cells. Thus, it is unlikely that electrical coupling or disynaptic inhibition accounts for the rapid CS-induced inhibition of SSs in neighboring PCs.

**Figure 2.**
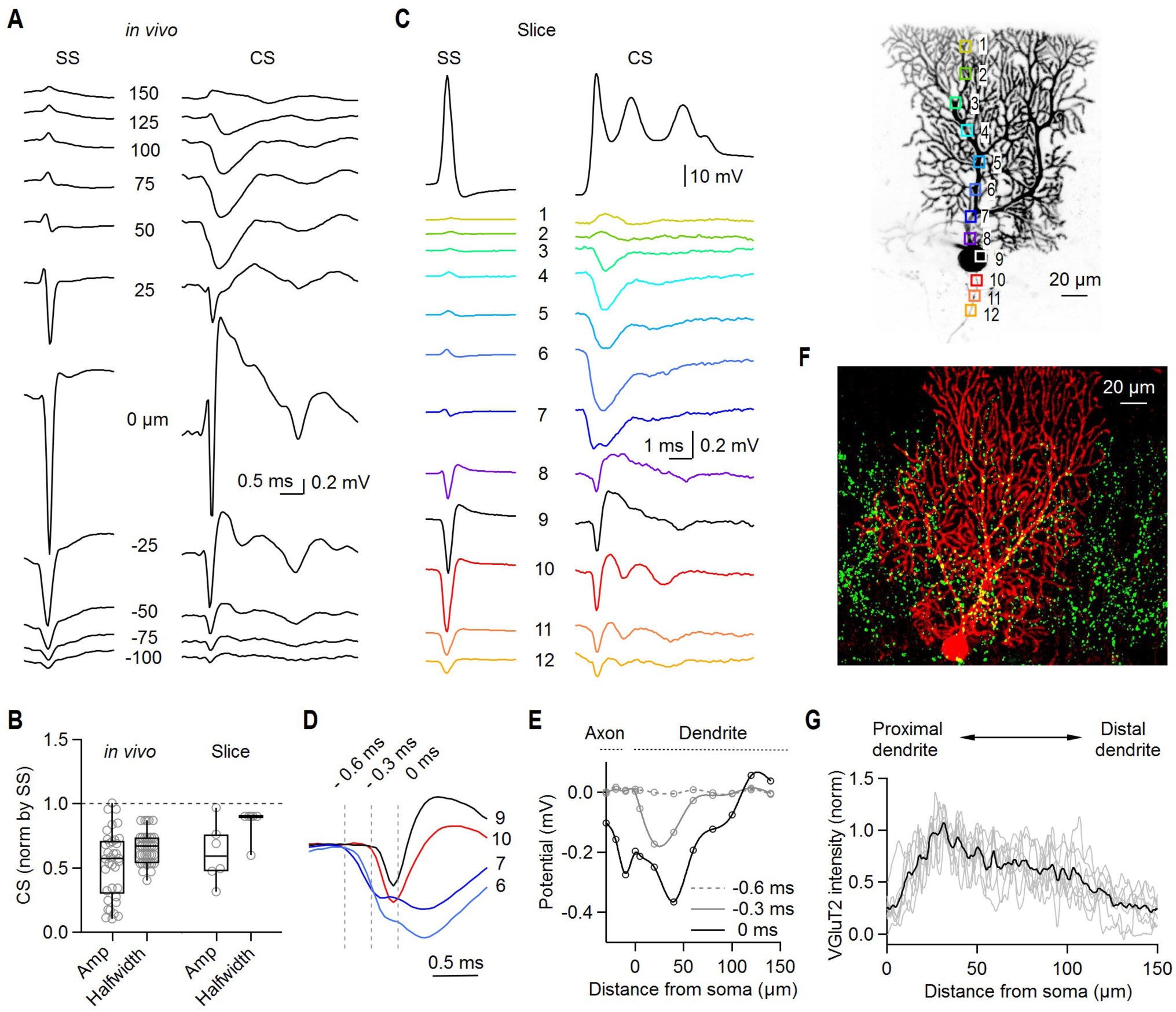
Extracellular potentials generated by simple spikes and complex spikes from Purkinje Cells *in vivo* and in brain slice. (A) Extracellular potentials recorded with a linear multielectrode array in a head-fixed awake mouse are shown for spontaneous simple spikes (SS) and complex spikes (CS). (B) Amplitudes and halfwidths of extracellular CS were normalized by SS properties for *in vivo* and slice recordings. (C) (top) SS (left) and CS (middle) responses measured by whole cell patch clamp in acute cerebellar slice for the fluorescently labeled cell (right). (bottom) Extracellular signals produced by SSs and CSs are color-coded, and were measured in the indicated regions dendrite (1-8), soma (9), and axon (10-12). (D) Average extracellular voltage changes produced by a CS measured in (C) are superimposed for the indicated region to compare their time course and amplitude (n=8 cells). (E) Summary of the amplitudes of extracellular potentials produced by a CS as a function of distance from soma at different time point indicated in (D) (n=8 cells). (F) Representative image of a PC (biocytin, red) in a slice immunostained for VGluT2 (green). (G) Summary of VGluT2 immunostaining as a function of distance from the PC layer (average black, individual slices (grey) (n=8 slices).

We next considered ephaptic coupling as a possible mechanism for CS inhibition of SSs. We have previously shown that ephaptic coupling allows PCs to rapidly signal to neighboring PCs to promote synchronous firing (Han et al., 2018). We found that for the cells in our study, SS activity for PC pairs was synchronized and that for a pair of PCs, the larger the extent of SS synchronization the more strongly CSs in one PC inhibit SSs in the other PC (Figure S4). It is surprising that CSs in one PC suppress SSs in neighboring cells, whereas SSs in the same PC elevate SS spiking in nearby PCs (Han et al., 2018). How can CSs and SSs have inversely correlated effects on the firing of neighboring cells? Is it possible that ephaptic coupling allows CSs to suppress and SSs to promote spiking in neighboring PCs?

It is expected that SSs and CSs produce very different extracellular signals. For SSs, sodium influx near the axon initial segment (AIS) of one cell produces a local hyperpolarizing extracellular potential large enough to activate sodium channels in the axons of neighboring PCs (by producing a depolarization of the net transmembrane voltage of the AIS). The extracellular signals associated with a CS are much more complicated. CF activation initially opens AMPA receptors, depolarizing the PC dendrite, cell body, and AIS and triggering dendritic calcium entry, evoking a short-latency sodium spike followed by a series of smaller spikelets, and ultimately activating potassium channels. The extracellular voltage changes evoked by CF stimulation thus result from a combination of current flow from AMPA receptors and multiple voltage-dependent channels on dendrites, cell body, and the axon. Consequently, the spatiotemporal extracellular signals are expected to be complex.

As a first step in evaluating possible ephaptic signaling by CSs, we measured the associated extracellular voltage signals at different regions of PCs *in vivo*. We used a linear electrode array oriented to record from dendritic, somatic, and axonal compartments of a PC (in contrast to Figure 1 in which we oriented the electrode parallel to the PC layer to record from multiple PCs). In these recordings we also measured extracellular signals associated with SSs (Figure 2A, left), which was useful in determining the location of the AIS. As shown in this recording, the site at 0 μm shows SS and CS signals similar to what has been reported previously (Bell and Kawasaki, 1972; Bosman et al., 2010; de Solages et al., 2008; Ebner and Bloedel, 1981; Wise et al., 2010). The initial sodium spike component of the CS is narrower and has a smaller absolute amplitude than a SS and is preceded by an initial positive signal (Figure 2B). These factors may limit the ability of the sodium spike component of the CS to mediate ephaptic excitation of PCs.

The signals in the dendritic region are very different for the CS than for the SS. SS signals are strongly attenuated 50 μm from the AIS because most of the current flow associated with a sodium spike occurs in the AIS. For a CS, we observed a hyperpolarized extracellular potential throughout the proximal dendrite (50-125 microns) that flipped in sign at the surface of the molecular layer and near the soma. This is consistent with current flow into the proximal dendrites and out of the distal dendrites and soma. Qualitatively similar results were obtained for 6 cells. However, a drawback of *in vivo* recordings of extracellular signals is that it is very difficult to know the precise position of electrode sites relative to the cell of interest.

We therefore complemented the *in vivo* recordings by measuring signals in acute brain slices using fluorescent dyes in PCs to precisely guide electrode placement (Figures 2C-2E). We obtained whole-cell recordings from PCs and measured spontaneous SSs and CSs evoked with a bipolar extracellular electrode placed in the granular layer or in the white matter (Figure 2C, top). Selective CF activation was achieved as described (Figure S5). We visualized PCs by including an Alexa dye in the recording electrode and measured extracellular signals from the indicated regions using a second electrode (Figure 2C, right). Extracellular signals evoked by SSs and CSs in slice were qualitatively similar to those recorded *in vivo*, with a prominent fEPSP extending 25 μm to 100 μm from the soma (Figure 2C middle, blue traces 3-7). Like the sodium spikes *in vivo*, evoked CSs in slice also had a smaller and narrower extracellular negativity relative to the SS (Figure 2B). The amplitudes of CSs and SSs were smaller in slice recordings than *in vivo*, as reported previously for SSs (Han et al., 2018). This is consistent with the extracellular solution shunting signals in slice recordings.

To determine the spatial origin of depolarizations contributing to the CS waveforms, we examined the timing of the extracellular signals with respect to the CF input. Extracellular signals near the proximal dendrite (regions 6 and 7) preceded signals near the soma and the AISs (regions 9 and 10) (Figure 2D). This is also illustrated by a summary of the amplitudes of extracellular signals recorded at different times (Figure 2E). These findings are consistent with the initial signal arising from inward current through AMPA receptors into dendrites (−0.6 ms). At 0 ms there is also a large inward current at the soma, which is consistent with a synaptically-driven sodium spike. This dendritic extracellular signal is well suited to rapidly influence nearby PCs.

These findings suggest that the location of CF synapses could be an important determinant of the site of the initial extracellular signals. We labelled individual PCs with biocytin and stained for CF boutons with an antibody to VGluT2 (Fremeau et al., 2001). CF boutons are most abundant in proximal dendrites 30-50 μm from the soma, and the density of boutons decreases towards the top of the molecular layer (Figures 2G), consistent with pervious results (Fremeau et al., 2001; Hioki et al., 2003; Miyazaki et al., 2003). This is apparent in the image where the PC dendrites extend beyond the regions containing CF boutons (Figure 2F). These findings (Figures 2F and 2G) suggest that the high CF bouton density in the proximal dendrites accounts for the rapid fEPSPs seen in Figures 2A and 2C (50 μm and region 6 respectively), and the lack of boutons in the distal dendrites helps to explain the positive signals at the top of the molecular layer (Figure 2A, 150 μm and Figure 2C, regions 1-2).

To gain insight into the mechanism of CS inhibition of SSs in nearby cells, we recorded from two neighboring PCs in voltage-clamp and stimulated the CF in one of the PCs (Figure 3A). The GABA_A_R antagonist GABAzine was included to exclude synaptic inhibition by interneurons. We stimulated extracellularly at an intensity that stochastically activated the CF input to one of the cells (PC1) on approximately half of the trials. We then divided PC1 responses into successes and failures and subtracted the average success from the average failure (Figure 3B, top). We used a similar procedure to isolate the currents in PC2 associated with activation of a CF synapse onto PC1 (Figure 3B, bottom). We found that the CF-triggered EPSC from one PC evoked rapid short-latency outward currents from nearby PCs (Figures 3B and 3C). This outward current is ideally suited to rapidly suppress SS activity.

**Figure 3.**
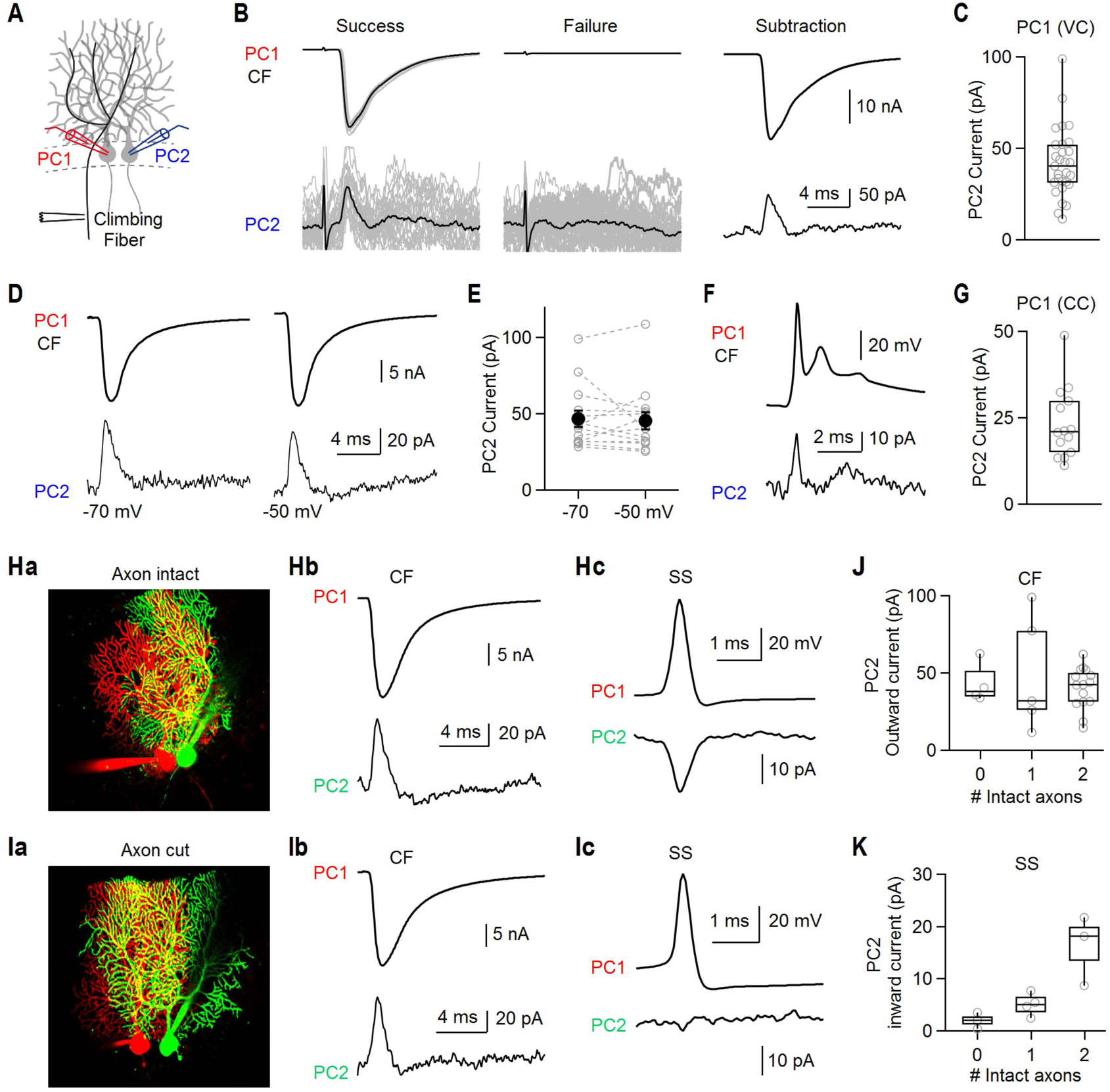
Climbing fiber stimulation evokes outward current in nearby Purkinje Cells regardless of whether axons are intact. (A) Schematic of experimental configuration with whole-cell recordings from two neighboring PCs, and a stimulus electrode positioned to activate selectively a CF innervating PC1. (B, top) Whole cell CF current measured from PC1 in voltage clamp mode at -70 mV. A threshold stimulus intensity was used that stochastically evoked successes (left) or failures (middle) (individual trials: grey and average: black). Average success – average failure is shown (right) (B, bottom) Simultaneous recordings from a nearby PC2 showed that successful stimulation of the CF input to PC1 was accompanied by an outward current in PC2 (left), but no such current was observed in PC2 when stimulation failed to evoke a CF input to PC1 (middle). Average success – average failure is shown (right). (C) Summary of outward current amplitudes recorded from PC2 when the CF innervating PC1 was activated and PC1 was voltage clamped at -70 mV. (D) The CF EPSCs in PC1 (HP=-70 mV, top), and currents recorded from nearby PC2 voltage clamped at either -70 mV (bottom left) or -50 mV (bottom right). (E) Summary of current amplitudes for PC2 at holding potentials of -70 mV and -50 mV. Paired t-test (p=0.69). (F) (top) Complex spike from PC1 recorded in current clamp. (bottom) Outward current from PC2 voltage clamped at -70 mV. (G) Summary of currents from PC2 accompanying complex spikes recorded in PC1 in current clamp (CC). (H, a) Representative image showing PC pair with intact axons. (H, b) CF-EPSC measured in in PC1 in whole cell voltage clamp (top) and current from nearby PC2 measured in voltage clamp (HP=-70 mV). (H, c) Spontaneous simple spike (SS) measured from PC1 in current clamp (top), and spike-triggered average current from PC2 in voltage clamp at -50 mV (bottom). (I) Same as (H) but with two cut axons. (J) Summary of the magnitudes of outward currents in neurons as a function of the number of intact axons for climbing fiber EPSC in neighboring cells (PC2 HP=-70 mV). (K) Same as (J) but inward currents by simple spikes (PC2 HP=-50 mV).

In order to clarify the currents evoked in neighboring PCs, it is important to determine the voltage dependence. Previously, we showed that SSs in one PC promote firing in neighboring PC pairs by evoking an inward current in neighboring cells. This current is mediated mainly by voltage-dependent sodium channels and is consequently steeply voltage-dependent, being much larger at -50 mV than at -70 mV (Han et al., 2018). We therefore compared the outward current evoked by CFs in neighboring PCs by voltage clamping PC2 at either -50 mV or -70 mV. In contrast to the situation with PC stimulation, the outward currents in PCs evoked by CF stimulation were not voltage dependent (Figures 3D and 3E). This suggests that CF inputs in one PC induce passive outward currents from nearby PCs.

We next performed experiments under more physiological conditions by recording from PC1 in current clamp. In this configuration, the CF-elicited complex spike of PC1 also evoked an outward current in nearby PC2 that had a prominent transient short latency component, as well as some smaller longer lasting components (Figures 3F and 3G). Outward currents in neighboring PCs were smaller when PC1 was in current clamp as opposed to voltage clamp. This decreased amplitude in current clamp compared to voltage clamp is consistent with the dendritic depolarization in current clamp lowering the driving force and reducing the amplitudes of synaptic currents.

We have previously shown that ephaptic coupling of SSs in nearby PCs requires intact axons. We therefore directly assessed the involvement of axon-to-axon communication in CS suppression of SSs in neighboring cells. Whole-cell recordings were obtained from PC pairs, and cells were fluorescently labeled to determine if their axons were intact (Figures 3Ha and 3Ia). Both PCs were held at -70 mV in voltage clamp mode, and a CF input onto PC1 was activated. This evoked outward currents in neighboring PCs regardless of the number of intact axons (Figures 3Hb and 3Ib). We also examined SS signaling by allowing PC1 to fire spontaneously in current clamp and recorded the spike-triggered average (STA) from the other PC in voltage clamp at -50 mV. PC pairs with severed axons showed extremely small STA responses, whereas STA inward currents were prominent for PC pairs with two intact axons (Figures 3Hc and 3Ic). Thus, intact axons are required for STA inward currents generated by SSs (Figure 3K) but are not required for CF-induced outward currents (Figure 3J). These findings establish that CF activation produces outward currents in neighboring cells through a mechanism that is independent of axons.

To test if AMPAR activation during CF stimulation is responsible for the outward current observed in nearby PCs, we recorded from two neighboring PCs and stimulated the CF in one of the PCs in the presence of NBQX. NBQX treatment completely blocked both the CF stimulation-elicited EPSC and the outward current from nearby PCs (Figures 4A and 4B). Application of cyclothiazide, a drug that prevents AMPAR desensitization, increased the AMPA response during CF stimulation and increased the outward current in neighboring PCs (Figures 4C and 4D). Based on the experiments thus far, we hypothesize that AMPA receptor-mediated synaptic currents flowing into PCs produce a negative extracellular signal that in turn evokes a passive outward current in neighboring PCs. If this is the case, we should be able to reverse the extracellular signal and the resulting current in the second PC by voltage clamping the first PC at positive potentials where AMPA receptor currents would be reversed. To test this prediction, we obtained whole-cell recordings from PCs using pipettes containing CsF-based internal solution with QX314 to block Na^+^ channels, and recorded extracellular signals at the indicated locations (Figure 4E). This approach allowed us to isolate the synaptic component associated with CF activation free from the complication of sodium spikes in the PC. Large negative extracellular signals critical for the inhibition of nearby cells were detected near proximal dendrites (Figure 4F, left, Figure 4G). Changing the holding potential from -70 mV to + 30 mV reversed the direction of the extracellular signals (Figure 4F, right, Figure 4G). We went on to determine whether this also altered the direction of currents evoked in neighboring PCs. In this case, we recorded from both PCs using pipettes containing CsF-based internal solution with QX314 (Figure 4H). When both PCs were voltage-clamped at -70 mV, the CF-triggered outward current was still observed in PC2 (Figure 4I, left, Figure 4J). This suggests that voltage-dependent Na^+^ or K^+^ channels are not required for an outward current. When the holding potential of PC1 was changed to +30 mV, the direction of whole cell current from PC1 was reversed, and the current from nearby PC2 was also reversed (Figure 4I, right, Figure 4J). These observations are consistent with the extracellular voltage change producing a passive current in neighboring PCs with a direction dictated by the direction of the extracellular signal.

**Figure 4.**
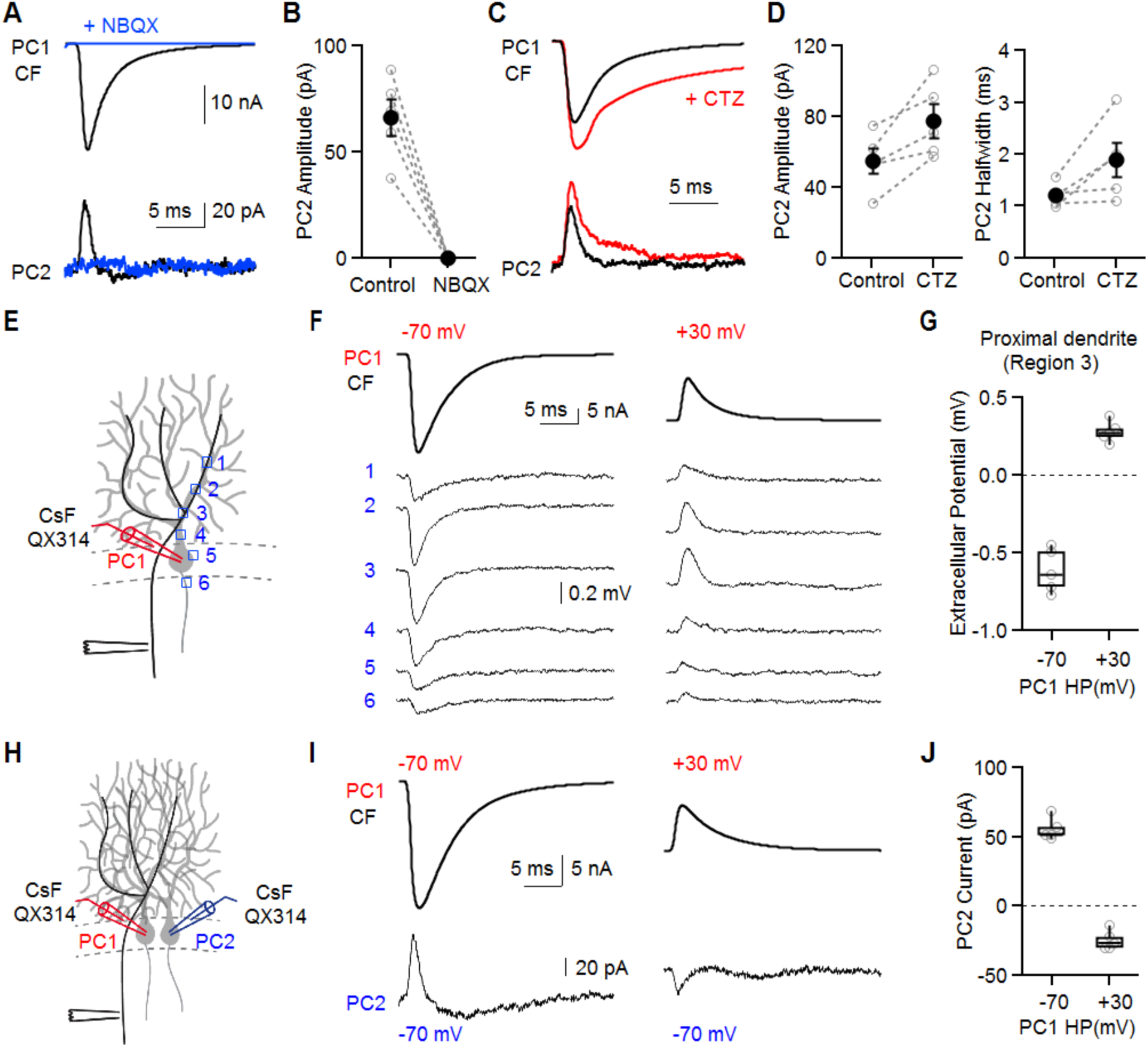
Reversing the climbing fiber EPSC by AMPARs activation in one Purkinje cell reverses extracellular currents and currents in neighboring cells. (A) The CF EPSCs in PC1 and currents recorded from nearby PC2 before (black) and after NBQX (5 μM, blue). (B) Summary of current amplitudes for PC2. (C) The CF EPSCs in PC1 and currents recorded from nearby PC2 before (black) and after cyclothiazide (100 μM CTZ, red). (D) Summary of current amplitude (left) and halfwidth (right) for PC2. (E) Schematic showing climbing fiber stimulation, whole cell voltage clamp recording and extracellular recording sites. (F) (top) Whole cell current from PC1 in voltage clamp at -70 mV (left) and +30 mV (right). (bottom) Extracellular voltage change was measured at dendrite (1-4), soma (5), and axon initial segment (6) indicated in (E). (G) Summary of the extracellular potential at the proximal dendrite (region 3). (H) Schematic showing paired recording with cesium fluoride (CsF) internal solution supplemented with the intracellular sodium channel antagonist QX314 (1 mM). (I) Climbing fiber EPSC from PC1 in voltage clamp at -70 mV (left) and +30 mV (right), and corresponding whole cell currents from nearby PC2 (HP=-70 mV). (J) Summary of the effects of the holding potential in PC1 on the currents measured in PC2 following climbing fiber activation of PC1.

Here we see that negative extracellular signals near the dendrite evoke an outward current in PCs, whereas we previously found that negative extracellular signals near the AIS evoke inward currents (Han et al., 2018). To clarify the discrepancy between axonal and dendritic stimulation, we used an extracellular electrode to directly locally alter the extracellular voltage either near the axon or the dendrites (Figure 5A), and we measured the resulting whole-cell currents and changes in SS firing in nearby PCs. The waveform for extracellular stimulation was a Gaussian curve with a half width of 1.2 ms (Figure 5B, top). Depolarizing extracellular stimulation near the dendrite induced an inward current in neighboring PCs, and hyperpolarizing stimulation induced an outward current in neighboring PCs. The amplitude of the evoked current was linearly related to the stimulus amplitude as expected for a passive current (Figure 5C, top). Signals evoked with dendritic stimulation were not influenced by the holding potential of the cell (Figures 5B and 5C). The same stimulation near the axon evoked very different currents at different holding potentials. At -70 mV, depolarizing extracellular stimulation evoked inward currents and hyperpolarizing extracellular stimulation evoked outward currents (Figure 5B, bottom blue, Figure 5C, bottom blue). These signals were scaled down versions of those evoked by dendritic stimulation. Holding potential had a large influence on currents evoked by extracellular stimulation near the axon. At -50 mV the direction of the intracellular signal was reversed (Figure 5B, bottom red, Figure 5C, bottom red). Differences between dendritic and axonal stimulation are consistent with dendritic signals being nearly passive because of the low density of Na channels. Axonal stimulation at a holding potential of -70 mV also evoked passive currents, but they were smaller because the surface area of dendrites is much larger than the surface area of the soma and axonal regions. The reversal in sign at -50 mV is a consequence of Na channel activation in the AIS due to the high density of Na channels.

**Figure 5.**
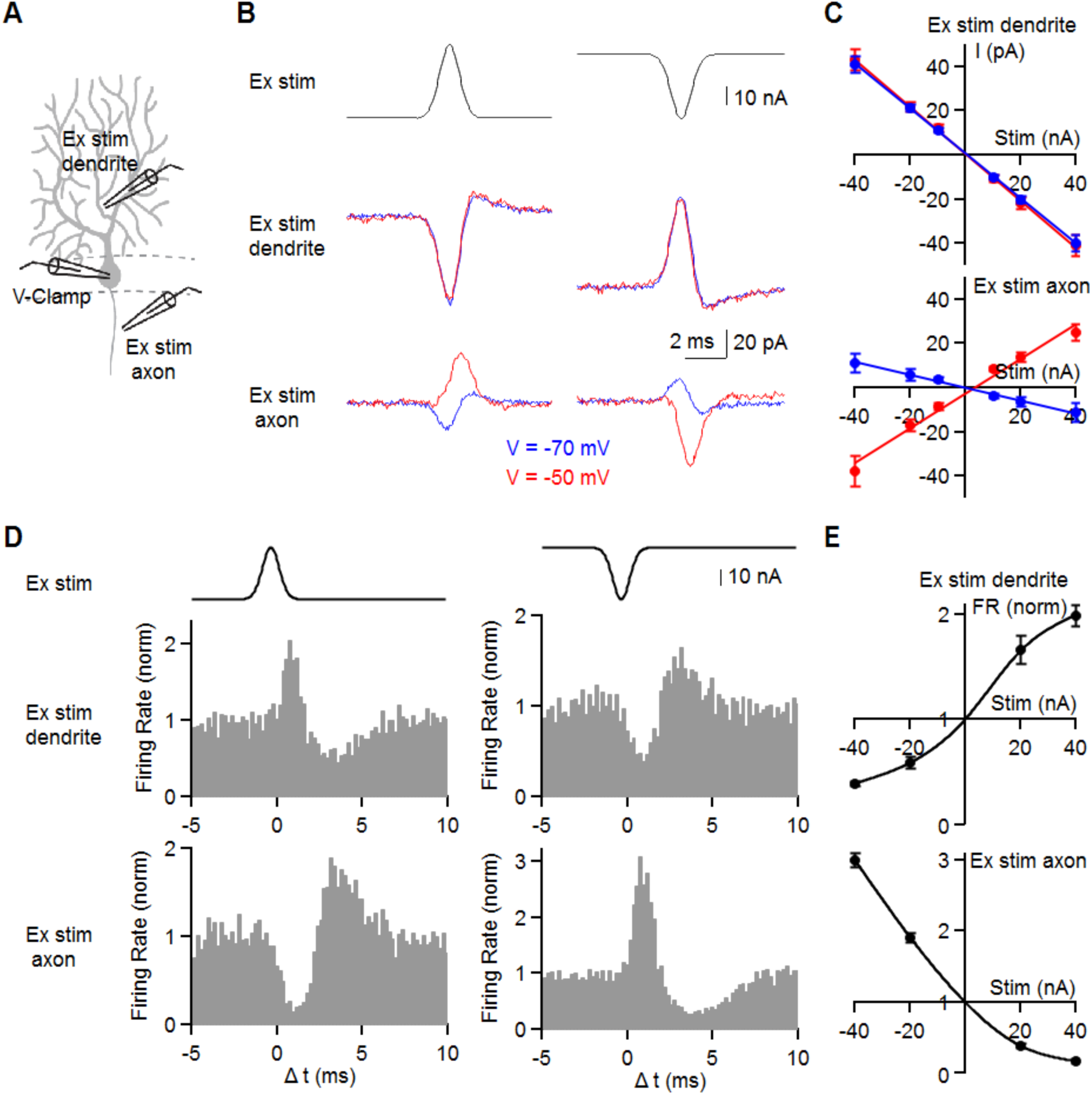
Differential effects of extracellular dendritic and axonal stimulation. (A) Schematic showing whole-cell voltage clamp recording from a PC and extracellular current injection near the dendrite (40 μm from soma) or near the axon (30 μm from soma) used in (B) and (C). In (D) and (E) the somatic recording is an on-cell recording to monitor SSs. (B) The waveform for extracellular stimulation (Ex stim, top) and the resulting currents measured at the holding potential of -70 mV (blue) and -50 mV (red) by extracellular stimulation near the dendrites or near the axon. (C) The relationship between the evoked currents and the intensity of extracellular stimulation near the dendrite (top) or axon (bottom) in voltage clamp at -70 mV (blue) and -50 mV (red). (D) Experiments similar to B but examining the effect of extracellular stimulation on spontaneous SSs measured with on-cell recordings. Histograms of firing rate in response to extracellular stimulation near the dendrites or the axon. (E) The relationship between firing rate and the intensity of extracellular stimulation near the dendrites (top) or the axon (bottom).

We then directly measured the effects of extracellular stimulation on spontaneous PC firing. We monitored spontaneous SS firing from PCs with an on-cell electrode and stimulated extracellularly with Gaussian waveforms either near the dendrites or the axon (Figures 5D and 5E). Depolarizing dendritic current stimulation rapidly elevated spiking, which was followed by a suppression of spiking (Figure 5D, middle left). Hyperpolarizing dendritic stimulation transiently suppressed spiking, which was followed by increased spiking (Figure 5D, middle right). In contrast, depolarizing current stimulation near the axon suppressed spiking and hyperpolarizing stimulation transiently elevated spiking (Figure 5D, bottom). Summaries of the influence of extracellular dendritic and axonal stimulation on PC firing rates illustrate the importance of stimulus position on the SS activity in a cell (Figure 5E). Thus, extracellular voltage changes near dendrites or axons exert opposite effects on PC firing.

We extended this approach to determine the effects of extracellular signals arising from a CS on a nearby cell. We separately stimulated the axon and the dendrites to assess the contributions of each of these aspects of the ephaptic signals that accompany a CS. We stimulated in 4 different ways using measured extracellular waveforms (Figure 6A). Extracellular stimulation of the axon with a waveform produced by a simple spike evoked an inward current in a PC clamped at -50 mV and a small outward current at -70 mV (Figure 6B). Stimulation with the extracellular signal produced by a CS near an axon evoked more complex currents with a much smaller inward component at -50 mV and small responses at -70 mV (Figure 6B). Stimulation with the extracellular waveforms produced by a CS near the dendrites evoked large inward currents that were very similar at -50 mV and -70 mV (Figure 6B). Stimulation of both axons and dendrites with CS waveforms evoked responses that are slightly larger and more temporally precise than those evoked by dendritic stimulation alone (Figure 6B).

**Figure 6.**
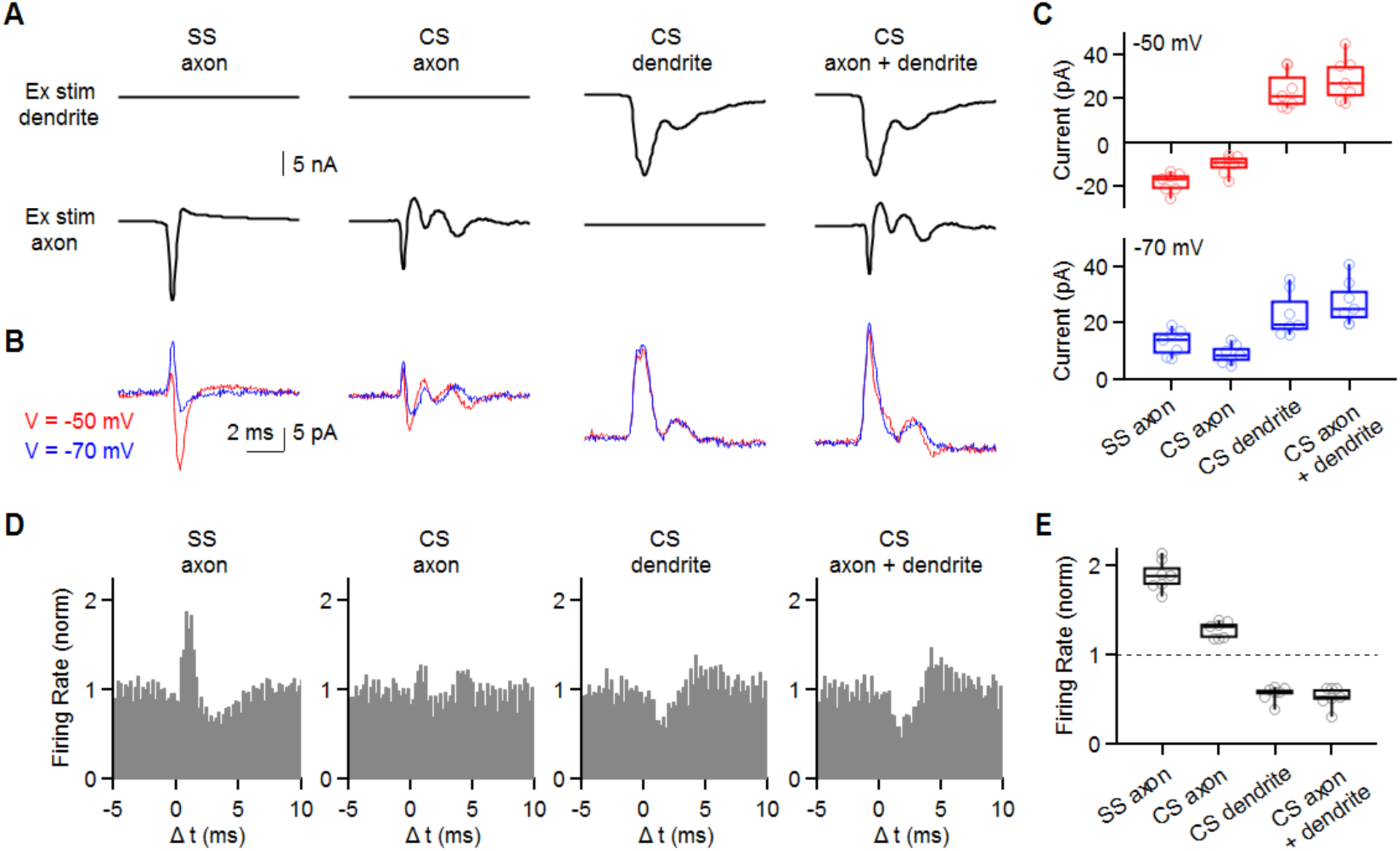
Mimicking extracellular signals that accompany a complex spike evokes outward currents and inhibits simple spikes. Experiments were performed similar to those in Fig. 5, but using waveforms based on extracellular signals accompanying complex spikes. (A) Stimulus waveforms corresponding to the waveform near the axon initial segment accompanying a SS (SS axon), the waveform near the axon initial segment associated with a complex spike (CS axon), the waveform near the dendrite associated with a complex spike (CS dendrite), and simultaneous axonal and dendritic stimulation with the appropriate CS waveform. Each waveform was injected into extracellular regions near the dendrite (top) and the axon (bottom). (B) Whole-cell currents evoked by the corresponding extracellular stimuli in (A) and recorded at a holding potential of either -70 mV (blue) or -50 mV (red). (C) (top) Summary of the current elicited by extracellular stimulation. (bottom) Same as (top) but -70 mV holding potential. (D) Histogram summaries simple spikes in response to the corresponding stimuli in A. (E) Summary of firing rate changes evoked by extracellular stimulation using the indicated waveforms.

**Figure 7.**
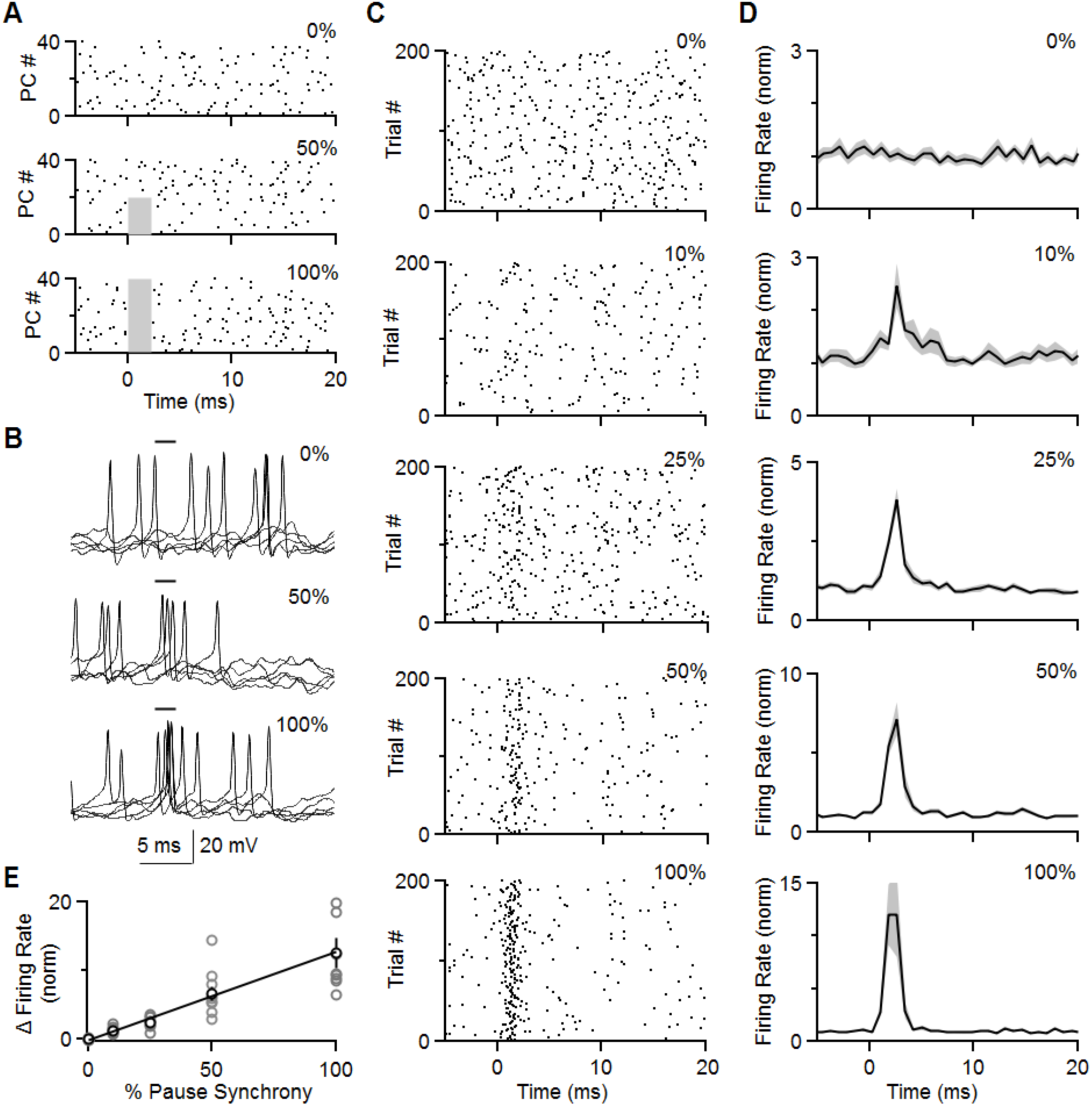
Dynamic clamp studies indicate that brief pauses in PC firing can effectively promote firing of neurons in the DCN. Dynamic clamp recordings were made from DCN neurons, with inhibition provided by 40 PC neurons with simple spike firing patterns as recorded *in vivo* (see Methods). PC spiking was suppressed for the indicated percentage of PCs, and the effect on DCN neuron spiking was determined. (A) Example spiking patterns are shown for 0 PCs for spikes eliminated for a 2 ms period (grey regions) in none of the PCs (top), half of the PCs (middle) and all of the PCs (bottom). (B) Examples show 10 superimposed traces for the same cell for no suppression (top), 50% of PCs (middle) and suppression in all PCs (bottom) for the two milliseconds indicated. (C) Raster plots for the same DCN neuron recorded when PC firing was suppressed for 2 ms in 0, 10, 25, 50, and 100% of PCs. (D) Summary of normalized histograms for DCN neurons when PC firing was suppressed for 2 ms in 0, 10, 25, 50, and 100% of PCs. (E) Summary of the effect of suppressing firing in a percentage of PCs on the normalized increase in DCN neuron firing frequency.

We then examined the effects of these stimuli (Figure 6A) on spontaneous SS firing (Figures 6D and 6E). The SS extracellular waveform evoked a large transient increase in spontaneous SS firing, but the CS axonal waveform had only a very small influence on firing (Figure 6D). Stimulation of the dendrites and simultaneous stimulation of the axon and dendrites with CS extracellular waveforms evoked a transient pause in SS firing that is similar to that observed *in vivo* (Figure 6D). In our slice experiments this brief pause is followed by a transient elevation in spiking that is not apparent *in vivo*. This may reflect our inability to perfectly mimic the extracellular signals evoked by a CS with just two electrodes.

### Functional implications

In order to evaluate the functional relevance of transient suppression of PC firing, it is necessary to consider the properties of PC outputs to neurons in the DCN. PC activity is conveyed to the rest of the brain primarily by regulating the activity of target neurons in the DCN. PCs are spontaneously active at high frequencies, and make strong inhibitory connections onto DCN neurons, with an estimated 40 PCs converging ono each DCN neuron (see Methods). Previous studies have raised the possibility that synchronous SS firing of a fraction of PCs can promote firing of DCN neurons (Han et al., 2018). Here we examine a related question: can a transient decrease in the firing of a fraction of PC inputs, as would result from CS-inhibition of SS firing in neighboring PCs, promote firing in the postsynaptic DCN neuron?

We first addressed this question by performing dynamic clamp experiments. The approach is similar to that taken previously to examine the consequences of synchronized PC spiking on DCN firing. Here we also assume there are 40 10nS inputs (Person and Raman, 2011). We based the presynaptic firing patterns on PC activity recorded in our *in vivo* experiments (Figure 1). We adjusted the amplitude of excitation (50-150 nS) to maintain the firing rate of DCN neurons in the 30 - 50 Hz range as recorded *in vivo* (Figure 8). We then measured the firing rate of DCN neurons when the firing of a fraction of the PC inputs was transiently suppressed for 2 ms (Figure 7A). Suppression of PC activity promoted short-latency spikes in DCN neurons (Figure 7B). As shown in raster plots for a representative DCN neuron, the transient suppression of PC firing promoted short-latency DCN neuron firing that became increasingly reliable as the percentage of suppressed PCs was increased (Figure 7C). A summary of the resulting histograms shows that, on average, a suppression of 10% (4 of 40 PCs) transiently increased DCN neuron firing by more than a factor of 2, and suppressing all of the PC activity increased DCN neuron firing approximately 10 fold (Figure 7D). There was an approximately linear relationship between the percentage of PCs suppressed and the increase in the DCN neuron firing (Figure 7E). It is clear that a mere 2 ms pause in firing of a fraction of PC inputs can rapidly and precisely control DCN firing.

**Figure 8.**
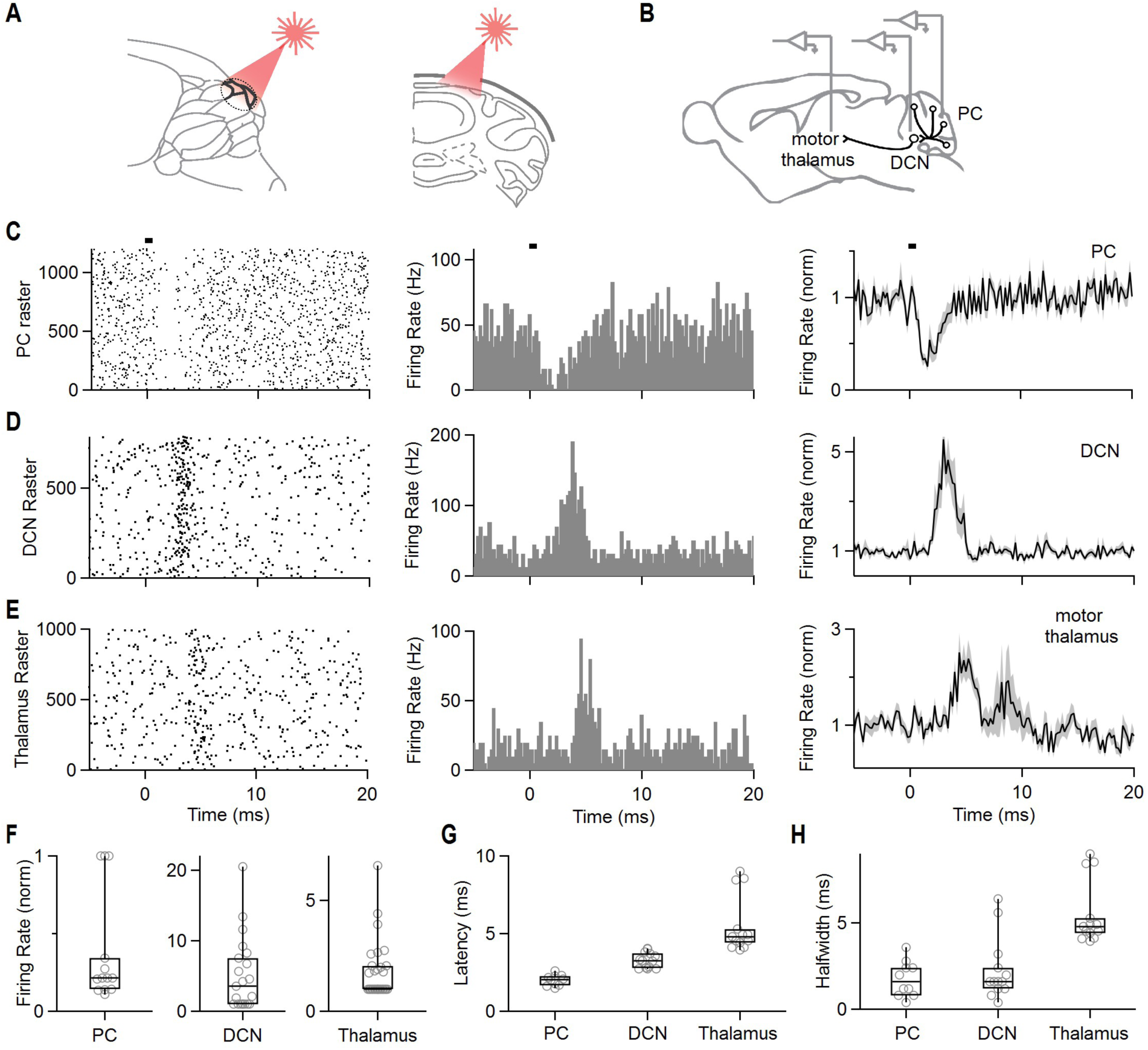
Brief suppression of Purkinje cell simple spikes increases firing in the DCN and thalamus. (A) Schematics showing craniotomy and illumination of a region of the cerebellar cortex. (B) Schematics showing *in vivo* recording in PC, DCN, and thalamus. (C) Raster plot of simple spikes from single PC (left). Histogram summarizing the data from left raster plot (middle). Average firing rate of simple spikes from PC by light stimulation (n=13). Shaded grey is SEM (right). Bar indicates 0.5 ms light stimulation at 0 ms. (D) Same as (C) but in the DCN. Average histogram is for all of the 21 cells in the DCN. (E) Same as (C) but in the thalamus (average histogram is for the 14 of 30 cells that responded to simulation). (F, G, H) Summary of normalized firing rates (F), latencies (G), and halfwidths (H) for PCs, DCN neurons, and thalamic neurons.

To test whether a brief pause in PC activity could also promote firing in the DCN *in vivo*, we used optogenetics to transiently suppress PC firing and recorded responses in DCN neurons. We exposed a large region of the cerebellar cortex (an approximately 1.5 mm diameter craniotomy) in a transgenic mouse that selectively expressed halorhodopsin in PCs, and suppressed PC firing with brief light flashes (0.5 ms, 80 mW, 647 nm see Method) (Figure 8A). We recorded the resulting activity in PCs, in DCN neurons, and in the motor thalamus (Figure 8B). This 0.5 ms stimulus suppressed PC firing for approximately 2 ms (Figure 8C), resulting in a time course similar to that arising from suppression induced by CSs in neighboring cells (Figure 1). Light suppressed the firing of most PCs we recorded from (regions directly below the craniotomy), although for some deep PCs suppression was not pronounced (Figure 8F). Because only a fraction of the total PCs in the cerebellum are close to the craniotomy, the majority of PCs are unlikely to be significantly suppressed because they will be exposed to much lower light levels. Nonetheless, our dynamic clamp studies suggest that simultaneous suppression of even a small fraction of PCs converging onto a DCN neuron will promote firing of that neuron. Firing was transiently elevated in all DCN neurons we recorded from (Figures 8D and 8F). Remarkably, 14 of 30 neurons also responded in the motor thalamus (Figures 8E and 8F), indicating that brief synchronous suppression of PCs not only drives DCN firing, it can also influence downstream regions targeted by the DCN (see also Figure S6). The latencies of the responses are consistent with rapid light evoked inhibition of PCs in ∼2 ms, followed ∼ 1 ms later by excitation of the DCN, and an additional 2.5 ms later in the motor thalamus (Figure 8G). The suppression of PCs was brief (halfwidth of ∼ 2 ms), as were the subsequent increases in spiking in the DCN (halfwidth of ∼ 2 ms), but the increases in the spiking of thalamic neurons was somewhat longer lived (halfwidth of ∼ 5 ms). These findings illustrate that in general brief synchronous pauses in PC firing can be highly effective in driving postsynaptic spiking. They also suggest that CS-induced pauses induced in PCs could be highly effective at influencing the spiking of neurons in the DCN.

## Discussion

Our main finding is that when a CF activates a Purkinje cell it simultaneously suppresses SSs in neighboring PCs. It is remarkable that a powerful glutamatergic excitatory synapse can rapidly and transiently inhibit spiking. We find that spiking is suppressed by a novel type of ephaptic signaling in which a single CF generates synaptic currents and extracellular signals of sufficient size to directly hyperpolarize neighboring cells.

### CSs do not promote SS firing in neighboring PCs

The observation that axonal SSs in one PC promote SS firing in neighboring PCs (Han et al., 2018), combined with the fact that CSs trigger a series of sodium spikes, raised the possibility that CSs could promote SS firing in neighboring cells. We found that this was not the case. Our analysis showed that two factors prevent a CS from stimulating neighboring PCs. First, the extracellular waveform near the initial segment is shorter-lived and has a lower peak amplitude than the extracellular signal associated with a SS. Consequently, it is not very effective at opening Na channels in nearby axons. Second, the negative signal in the dendrites that rapidly suppresses SSs precedes the excitatory signal near the axons. Consequently, the inhibitory effects dominate to suppress spiking in nearby PCs.

### The influence of location on effects of ephaptic signals

We found that the location of extracellular voltage changes from a stimulus electrode had a profound influence on the consequences of stimulation. For extracellular dendritic stimulation, positive stimuli promoted SSs and negative stimuli suppressed SSs. The opposite was true for axonal stimulation in which positive stimuli suppressed SSs and negative stimuli promoted SSs. Extracellular stimulation combined with voltage clamp experiments provided an explanation for these findings. When PCs were voltage clamped at - 70 mV, a potential where very few sodium channels are open, then extracellular stimulation of either dendrites or axons evoked currents that were in the same direction: positive stimuli promoted inward currents and negative stimuli promoted outward currents. This is consistent with passive effects of extracellular stimulation. The signals were larger for dendritic stimulation, as expected from the large surface area of dendrites, making them well suited to detecting extracellular signals. At a holding potential of -50 mV, currents evoked by extracellular dendritic stimulation were unchanged, but axonal stimulation evoked large signals in the opposite direction. This is consistent with the involvement of voltage-gated sodium channels in the axon initial segment, and the fact that at -50 mV sodium channel opening is highly sensitive to membrane potential (Carter et al., 2012).

Previous studies of ephaptic signaling and PCs focused on the axon initial segment (Blot and Barbour, 2014; Han et al., 2018). Positive extracellular signals near the axon associated with activation of MLI pinceaus suppressed PC firing whereas negative extracellular signals produced by sodium influx into an axon opened sodium channels in nearby PC axons to promote firing. Here we see just the opposite, in which a negative signal in the dendrites suppresses PC firing by passively inducing an outward current.

### A novel type of ephaptic signaling

The CS induced SS inhibition we describe here represents a novel form of ephaptic coupling. Previously, it was shown that the collective extracellular signals associated with the activity of many cells leads to slow correlations in firing (Anastassiou et al., 2015; Anastassiou et al., 2011). It had also been shown that by opening voltage-gated channels, single cells can generate sufficiently large extracellular signals to influence the axons of target cells in order to influence their excitability (Blot and Barbour, 2014; Furukawa and Furshpan, 1963; Han et al., 2018; Korn and Axelrad, 1980; Korn and Faber, 1975). The ephaptic signaling we describe here differs in that it reflects primarily the current flow through ionotropic channels of a single synapse consisting of hundreds of release sites. There are several reasons why this type of signaling is particularly effective for CSs in PCs. First, CF synapses are extremely powerful. Second, the dendritic arbors of PCs are very large, which make them well suited to detect the widespread signals generated by nearby PCs. Third, the PC dendrites are located in close proximity to each other. All of these factors combine to make CS to SS signaling particularly effective.

### Spatial extent of ephaptic inhibition

Our findings suggest that ephaptic signaling allows each IO neuron to simultaneously suppress firing in many PCs. An IO neuron gives rise to an average of 7 CFs onto different PCs (Sugihara et al., 2001), and whenever an IO neuron fires it evokes CSs in these target cells. Here we have shown that CSs inhibit SS firing in neighboring PCs up to 50 μm away. This suggests that each CS ephaptically inhibits SSs in a cluster of up to 20 PCs, and therefore a single IO neuron will simultaneously suppress firing SSs in approximately 100 PCs, based on the PC packing density (Doulazmi et al., 1999; Hadj-Sahraoui et al., 1996; Woodruff-Pak, 2006). This leads to synchronous disinhibition and effective excitation of downstream DCN neurons. Our findings suggest that the net effect of IO synapses in the cerebellar cortex might be to provide less overall inhibition to the DCN as a consequence of inhibiting the firing of so many PCs. Previous studies have observed excitation in the DCN in response to complex spikes in the PC layer (Blenkinsop and Lang, 2011). This has been attributed to CF collaterals that impinge on the DCN directly (De Zeeuw et al., 1997; Sugihara et al., 1999; Sugihara et al., 2001; van der Want and Voogd, 1987). The CS induced pause in the firing of nearby PCs we describe here could also contribute to increased responses in the DCN. In addition, *in vivo* recordings indicate that different IO cells can fire synchronously (Armstrong et al., 1968; Bell and Kawasaki, 1972; Crill, 1970; Welsh et al., 1995), which could greatly increase the number of PCs that synchronously pause. This suggests that a small number of IO neurons could have a profound impact on inhibition within the DCN that could be magnified if a large number of paused PCs converge onto the same population of DCN neurons.

### Functional significance of pauses in PC firing

Our dynamic clamp studies and optogenetic suppression studies establish that transient synchronous suppression of PC firing is remarkably effective at elevating the firing of DCN neurons. One of the striking features of this inhibition is that it leads to such rapid and precisely timed spiking in DCN neurons. Just one millisecond after the commencement of a pause in PC firing there is a large increase in the firing of DCN neurons (Figure 8). This is much more rapid than the stimulus to spike latency of 5-10 ms that is observed following synchronous IPSCs (Person and Raman, 2011). One of the most remarkable and poorly understood features of the cerebellum is that the output of the cerebellar cortex relies on inhibitory neurons that are spontaneously active at very high frequencies. This means that for a DCN neuron receiving inputs from 40 PCs firing at 100 Hz, the total frequency of inhibitory inputs is on the order of 4 kHz. This is an important feature that allows decreases in firing to rapidly promote firing in DCN neurons. The percentage of PCs that synchronously pause in their firing can strongly influence the response of neurons in the DCN. Our dynamic clamp studies suggest that a pause in spiking in just 10% of the PC inputs (4 of 40 inputs), will lead to an approximate doubling of DCN neuron spiking. Based on the large number of PCs expected to be inhibited by a single spike in an IO neuron, this suggests that a single IO neuron might be able to promote firing in many DCN neurons by suppressing firing of PC neurons.

## STAR Methods

### Key Resource Table

| Reagent or Resource | Source | Identifier |
| --- | --- | --- |
| **Antibodies** |  |  |
| Streptavidin, Alexa 568 | Thermo Fisher Scientific | Cat# S11226 |
| Guinea Pig anti-VGluT2 | Synaptic Systems | Cat# 135 404 |
| Alexa 488 goat anti-guinea pig IgG | Abcam | Cat# ab150185 |
| **Chemicals and Peptides, and Recombinant Proteins** |  |  |
| SR95531 (Gabazine) | Abcam | Cat# ab120042 |
| QX-314 Chloride | Abcam | Cat# ab120118 |
| Alexa 488/594 hydrazide | Thermo Fisher Scientific | Cat# A10436/A10438 |
| Biocytin | Sigma-Aldrich | Cat# B4261 |
| Cyclothiazide | Abcam | Cat# ab120061 |
| NBQX | Abcam | Cat# ab120046 |
| Triton X-100 | Sigma-Aldrich | Cat# T8787 |
| R-CPP | Abcam | Cat# ab120159 |
| Strychnine hydrochloride | Sigma-Aldrich | Cat# S8753-25G |
| **Software and Algorithms** |  |  |
| Igor Pro 6 | Wavemetrics | https://www.wavemetrics.com |
| MafPC | Courtesy of M. A. Xu-Friedman | https://www.xufriedman.org/mafpc |
| Offline sorter | Plexon | https://plexon.com/products/offline-sorter/ |
| Matlab (R2017a) | Mathworks | https://www.mathworks.com/products/matlab |
| ImageJ | NIH | https://imagej.nih.gov/ij/ |

### Contact for Reagent and Resource Sharing

Further information and requests for resources and reagents should be directed to and will be fulfilled by the Lead Contact, Wade G. Regehr.

### Experimental Model and Subject Details

#### Mice

All animal experiments were performed according to NIH guidelines and protocols approved by Harvard University. P30-P45 C57BL/6 mice (Charles River) of either sex were used for acute slice experiments, and P50-P90 C57BL/6 mice (Charles River) of either sex used for *in vivo* experiments. We used Pcp2-cre mice (Jackson Laboratory, stock number 010536) crossed to eNpHR3.0-EYFP (Halo) mice (Ai39 Jackson Laboratory, 014539) for brief inhibition of Purkinje Cell activity *in vivo*. cKit-Cre mice (generous gift from J. Christie) were used for optogenetic inhibition of molecular layer interneurons.

## Method Details

### *In vivo* recording

*In vivo* recordings from PCs were made from awake, head-restrained mice using an acutely inserted silicon probe. To prepare for recordings, mice were anesthetized with 2% isoflurane and implanted with a head-restraint bracket using metabond (Parkell, Edgewood, NY). A craniotomy (∼ 0.5 mm diameter) was made over the vermis (centered on the midline-6.8 mm posterior to bregma). Craniotomies were sealed with kwiksil (World Precision Instruments, Sarasota, FL) until the day of the recording. After surgery, mice were given the analgesic buprenorphine. Mice were head-fixed over a cylindrical treadmill to allow for free locomotion. All animals were acclimated for at least one session prior to the recording session.

To measure the inhibition of PC firing after complex spikes in nearby cells, recordings were made using either the P or H2 style silicon probes from Cambridge Neurotech (Cambridge, England). The P style probe consists of two rows of 8 contacts, separated by approximately 25 μm (center to center distance). The H2 style consisted of a single row of 16 contacts, also separated by 25 μm. PCs were identified by their complex spikes. Care was taken to have clear, discernable, and individual spiking with minimal signal overlap in neighboring channels during simultaneous recordings. To clearly resolve the effect of complex spike activity on neighboring PCs, recordings lasted for at least 10 minutes. 6 of 37 PC pairs in Figure 1 were reanalyzed from our previous paper’s dataset (Han et al., 2018).

Exclusively H2 style probes with the linear arrangement of recording sites were used for measurements of extracellular action potential waveform shapes across the PC’s longitudinal axis (Figure 2A). To sample PCs along this direction, care was taken to insert the probe perpendicularly to the surface of the cerebellum, and thus, many of the underlying PC layers. When positioned correctly, the simple spike waveform transitions from a predominantly negative waveform near the axon initial segment to one with an initial prominent positivity at the soma/proximal primary dendrite. We used this characteristic transition across the linearly arranged recording sites to determine whether our probe was positioned perpendicular to the PC layer. Furthermore, the absence of significant PC activity in neighboring channels was also a clear indicator that only one contact on the probe was situated in the PC layer, and the others were above or below it. The PCs were positioned such that they were “centered” on the silicon probe, with at least 5 contacts above and 5 contacts below the central recording site.

### Acute slice preparation

Acute parasagittal cerebellar slices of 200∼250 μm thickness were prepared from the vermis. Mice were anesthetized with an intraperitoneal injection of a ketamine (100 mg/kg) and xylazine (10 mg/kg) mixture, and transcardially perfused with an ice-cold solution containing the following (in mM): 110 Choline Cl, 7 MgCl_2_, 2.5 KCl, 1.25 NaH_2_ PO_4_, 0.5 CaCl_2_, 25 Glucose, 11.6 Na-ascorbate, 2.4 Na-pyruvate, and 25 NaHCO_3_ equilibrated with 95% O_2_ and 5% CO_2_. The cerebellum was dissected, and slices were made using a VT1200s vibratome (Leica, Buffalo Grove, IL) in ice-cold solution as above. Slices were then transferred to a submerged chamber with artificial cerebral spinal fluid (ACSF) composed of (in mM): 125 NaCl, 26 NaHCO_3_, 1.25 NaH_2_PO_4_, 2.5 KCl, 1 MgCl_2_, 1.5 CaCl_2_, and 25 glucose, equilibrated with 95% O_2_ and 5% CO_2_, and allowed to recover at 32°C for 20 min before cooling to room temperature.

### Acute slice electrophysiology

Cerebellar slices were transferred to the recording chamber, and constantly perfused with ACSF containing the following (in mM): 125 NaCl, 26 NaHCO_3_, 1.25 NaH_2_ PO_4_, 2.5 KCl, 1 MgCl_2_, 1.5 CaCl_2_, and 25 glucose, equilibrated with 95% O_2_ and 5% CO_2_ at 32°C. PCs were visually identified with infrared differential interference optics. Patch pipettes of 1-3 MΩ resistance pulled from borosilicate capillary glass (Sutter Instrument, Novato, CA) with a Sutter P-97 horizontal puller. For whole-cell recordings, electrodes were filled with an internal solution containing (in mM): 150 K-Gluconate, 3 KCl, 10 HEPES, 0.5 EGTA, 3 Mg-ATP, 0.5 Na-GTP, 5 Phosphocreatine-tris_2_, and 5 Phosphocreatine-Na_2_ adjusted to pH 7.2 with KOH. The osmolarity was adjusted to 310 mOsm. The internal solution for Figure 5 contained (in mM): 100 CsCl, 35 CsF, 10 EGTA, 10 HEPES, and 1 QX314 adjusted to pH 7.2 with CsOH, and 310 mOsm. For on-cell recordings, glass pipettes were filled with ACSF. Electrophysiology data were acquired using a Multiclamp 700A amplifier (Axon Instruments, San Jose, CA) and an ITC-18 (Heka instruments Inc. Holliston, MA), filtered at 10 kHz, sampled at 20 kHz, and saved using software custom written in Igor Pro (Lake Oswego, OR).

### Biocytin labeling and Immunohistochemistry

Recordings of at least 20 minutes duration were obtained as described above, but with an internal solution supplemented with 0.2% biocytin. Slices were fixed overnight in 4% PFA at 4°C. Free-floating slices were then blocked for 4-6 hrs (5% Normal Goat Serum in 0.5% Triton X-100) at room temperature, followed by incubation overnight at 4°C with primary antibody (Guinea Pig anti-VGluT2, synaptic systems, 1:1000). Secondary antibodies (Streptavidin Alexa 568, Thermo Fisher Scientific, 1:1000; Alexa 488 goat anti-guinea pig IgG, Abcam, 1:1000) were then applied for 1 hr at room temperature. Slices were mounted using #1.5 coverslips and Prolong Diamond Antifade mounting medium. Z stacks (0.5 μm/section) were collected on an Olympus Fluoview1000 confocal microscope with FluoView software using a 60X 1.42 NA oil immersion objective.

### Purkinje cell labeling

Nearby PC pairs were whole-cell patched using an internal solution that included 50 μM of either Alexa-488 or Alexa-594 hydrazide (Thermo Fisher Scientific, Waltham, MA Alexa dye was allowed to diffuse into the cell for 10-15 min. Cells were then imaged with a custom built two-photon microscope, with image acquisition controlled by custom software written in Matlab (Mathworks, Natick, MA). Cells were imaged in 100-150 image planes and Z-spacing of 0.5 μm. Image contrast, brightness and Z-projection of images were processed in ImageJ (NIH, Bethesda, MA)

### Extracellular stimulation

The amplitudes and shape of extracellular current injection was based on recorded extracellular voltage responses of SS and CS measured at the axon initial segment and dendrite. The waveform was injected into extracellular space after convolution with a Poisson spike train (20 Hz) of unitary amplitude.

### Optogenetic Inhibition of Interneurons

CKit-Cre mice (generous gift from J. Christie) were used for in vivo inhibition of molecular layer interneurons. Mice were injected with 250 nl of AAV9-Ef1a-DIO eNpHR 3.0-EYFP (Addgene) at nine sites in the cerebellum and *in vivo* recordings were performed 2-3 weeks later. A 647 nm laser (Opto Engine, Midvale, UT) delivered via an 400 μm 0.39 NA optical fiber (Thorlabs, Newton, NJ) was focused to a spot of 2 mm^2^ centered over the recording site at an estimated power density of (40 mW/mm^2^). A silicon probe was lowered into the brain (P-series, Cambridge Neurotech, Cambridge, England), to find pairs of neighboring PCs within Halo-expressing regions. Once a pair was located, the posterior vermis was illuminated for 5 s on and 5s off for at least 30 minute. Recorded PC pairs were screened for an increase in activity in response to inhibition of MLIs. For analysis, complex spikes were sorted into laser on trials and laser off trials, and otherwise analyzed as previously discussed.

### Dynamic clamp

Sagittal slices of the DCN (180-200 μm) were cut from P30-P40 C57/BL6 mice of both sexes. Animals were anesthetized with ketamine / xylazine /acepromazine and transcardially perfused with warmed choline-ACSF solution containing (mM): 110 Choline Cl, 2.5 KCl, 1.25 NaH_2_PO_4_, 25 NaHCO_3_, 25 glucose, 0.5 CaCl_2_, 7 MgCl_2_, 3.1 Na-Pyruvate, 11.6 Na-Ascorbate, 0.005 NBQX, and 0.0025 (R)-CPP, oxygenated with 95% O2 / 5% CO2. Slices were transferred for 10-12 minutes to a standard ACSF solution (mM: 127 NaCl, 2.5 KCl, 1.25 NaH_2_PO_4_, 25 NaHCO_3_, 25 glucose, 1.5 CaCl_2_, and 1 MgCl_2_) and then allowed to rest at room temperature for 20-30 minutes before recordings. The same ACSF solution was used for recordings, with the addition of synaptic blockers (5 μM NBQX, 5 μM gabazine, 2.5 μM R-CPP, and 1 μM strychnine.) The same K-gluconate-based internal used for earlier current clamp experiments was used. Recordings were performed at 36°C.

Dynamic clamp recordings were made at 20 kHz with an ITC-18 computer interface (Heka, Holliston, MA) controlled by mafPC in Igor Pro (Wavemetrics, OR). 40 PC inputs with firing rates between 30 and 113 Hz based on our *in vivo* PC recordings. The number of PC inputs was chosen based on previous studies (Person and Raman, 2011; Wu and Raman, 2017). The intervals between spikes were randomized and convolved with a unitary synaptic conductance set at 3.7 nS conductance (10 nS inputs with a depression of 0.37 (Turecek et al, 2016) with a 0.1 ms rise time and a single exponential τ_decay_ of 2.4 ms (based on a fit to a recorded IPSC). 2 ms pauses of PC firing were administered every 250 ms before convolving with the unitary PC conductance. E_Inhibitory_ was set at -75 mV, accounting for a 7 mV liquid junction potential.

### Brief optogenetic inhibition of PC activity *in vivo*

Pcp2-cre mice crossed to eNpHR3.0-EYFP (Halo) mice were used to transiently inhibit PC activity *in vivo*. After preparing the mice for *in vivo* recordings, a craniotomy of approximately 1.5 mm diameter was made above the DCN. A 647 nm laser of approximately 2 mm in diameter was centered on the craniotomy. Steady state power density from the laser was measured to be 80 mW/mm^2^. Flashes of 0.5 ms were delivered to the region at approximately 2 sec intervals while recordings were made in the PCs above the targeted cerebellar nuclei, the nuclei themselves, or thalamic motor regions. PCs were identified by their characteristic firing rate, presence of complex spikes, and appearance in layers. DCN neurons were identified by depth, firing rate, and absence of complex spikes, Thalamic recordings were made targeting the motor subnuclei (ventrolateral aspect primarily) and confirmed by coating the recording electrode with DiI, followed by post-hoc histology. Recordings were made from head-restrained, awake mice, using E-series silicon probes (Cambridge Neurotech, Cambridge, England).

### Quantification and Statistical Analysis

#### Data analysis

Data analysis was performed using custom scripts in Matlab (Mathworks, Natick MA). Unpaired or paired Student’s t test were used when there were only two groups in the dataset. The statistical analysis used in each experiment, p-values, and the definitions of n are stated in the figure legends.

#### *In vivo* electrophysiology

Data were acquired at 20 or 30 kS/s (1 Hz to 10 or 15 kHz bandpass respectively) using an Intan acquisition system (Intantech). Spikes were sorted and aligned in Offline Sorter (Plexon). All further analysis and alignment between recordings (such that each complex spike associated timepoint was triggered on the same point on the complex spike waveform) was done in Matlab (Mathworks). Complex spikes from raw traces were subtracted for analysis of SS pause (Figure S1). Inhibition of simple spikes by the neighboring complex spike was quantified from neighboring complex spike-triggered peristimulus firing rate histograms: the average firing rate was taken between 0.5 and 1 ms after the complex spike and divided by the average firing rate of the cell (from the 100 ms prior to complex spike). Simple spike synchrony was calculated similar to reported previously (Han et al., 2018). Briefly, simple-spike triggered averages were normalized to the average event rate. A window between 0.6 ms before and after zero-lag was taken and averaged to determine the amount of synchrony.

#### Quantification of DCN and thalamic responses

To quantify the latency of responses in DCN and moto thalamus to PC inhibition, resulting firing rate PSTHs were baseline subtracted, and fit to a Gaussian function. Offset, amplitudes, and the width at half-max were calculated from the fitted data. Poorly fit responses (r^2^ < 0.2) were considered non-responders.

## Acknowledgments

This work was supported by National Institutes of Health Grant R35NS097284 to WR, an NIH postdoctoral fellowship F32NS101889 to C.H.C., the Stuart & Victoria Quan Fellowship in Neurobiology to C.G, and a National Science Foundation Graduate Research Fellowship under Grant No. (1745303) to M.M.K. We thank T. Osorno for the slice two-photon imaging. We thank B. Bean, L. Witter, and S. Rudolph for comments on the manuscript and thank Matthew Xu-Friedman for help with dynamic clamp.

## Author Contributions

K-S.H., C.H.C., and W.R. conceived the experiments. C.H.C. conducted most experiments in Figures 1, 8, S1, S2, S3, S4, and S6, MK did experiments in Figure 7, and K-S.H. conducted all other experiments. K-S.H., C.H.C., M.K. and C.G. did the analysis. K-S.H., C.H.C., and W.R. wrote the manuscript.

