## supplementary figures for "Climbing fiber synapses rapidly inhibit neighboring Purkinje cells via ephaptic coupling"

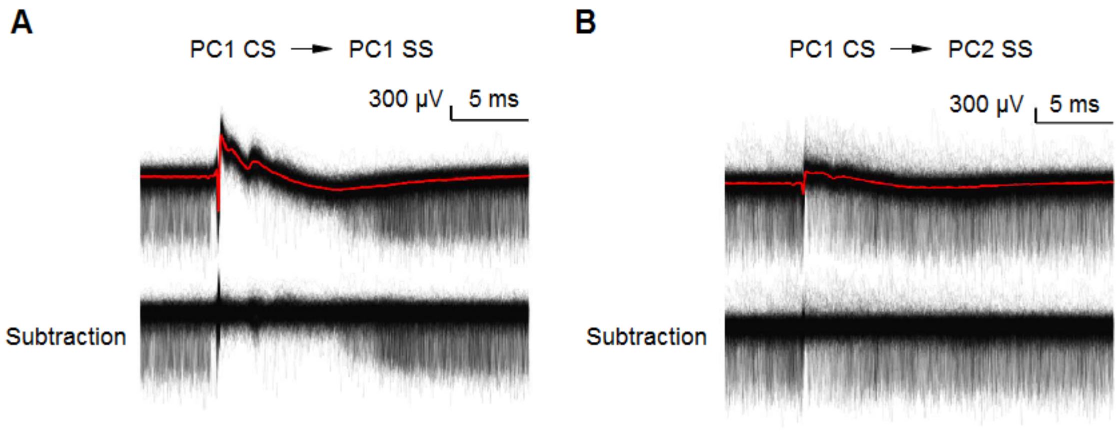


**Figure S1. Subtraction of complex spikes from raw traces, related to Figure 1.**
(A) (top) Simple spikes and complex spikes recorded on a same cell (PC1 SS) are aligned to PC1 CS (red). (bottom) Complex spikes from PC1 were subtracted.
(B) (top) Simple spikes simultaneously recorded on a neighboring site (PC2 SS) are aligned to PC1 CS. Complex spikes from PC1 (PC1 CS, red) were detected in neighboring site (PC2). (bottom) Complex spikes from PC1 were subtracted.

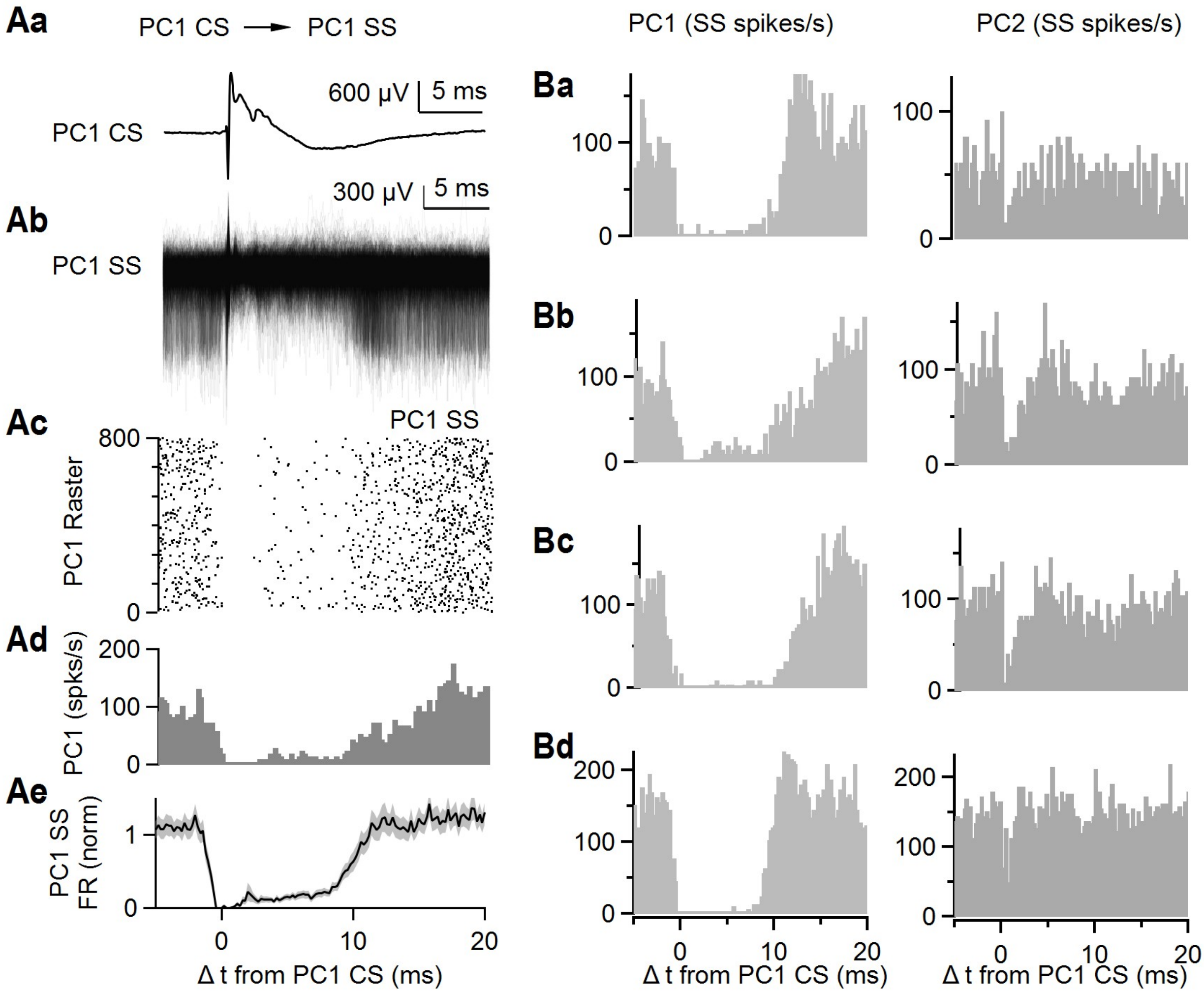


**Figure S2. Complex spikes in a Purkinje cell inhibit simple spikes in the same PCs in awake mice, related to Figure 1.**

(A, a) Average complex spike recorded on a single site (PC1 CS).
(A, b) Simple spikes simultaneously recorded on the same cell (PC1 SS) are aligned to PC1 CS.
(A, c) Raster plot of simple spikes from (A, b).
(A, d) Histogram summarizing the data in (A, b).
(A, e) Average of firing rate of simple spikes from the same PCs (PC1 SS) after complex spikes from PC1 (PC1 CS). Shaded grey is SEM.
(Ba-d) Four example pairs of nearest neighbor cells showing histograms of PC1 and PC2 SSs relative to CSs in PC1.

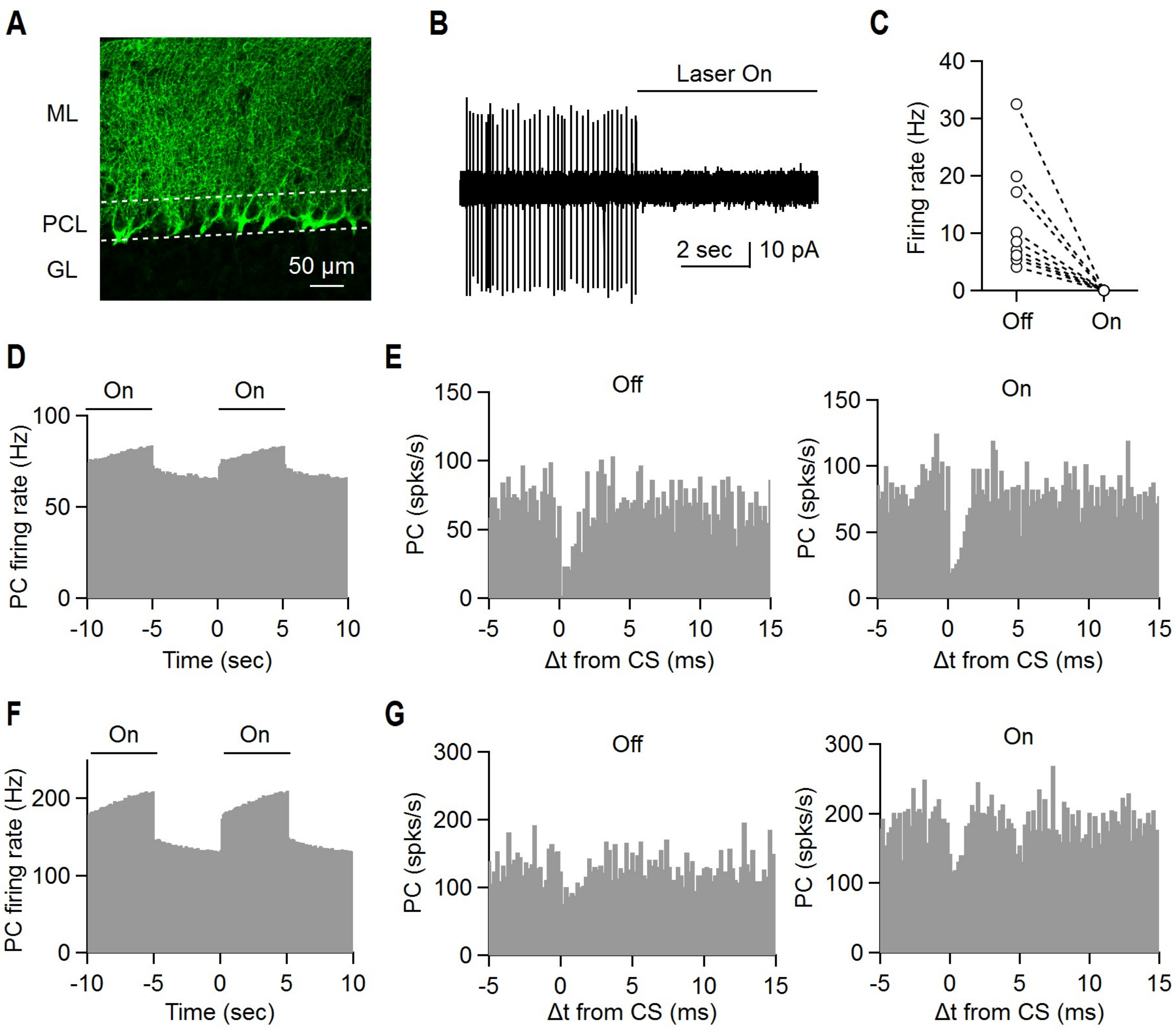


**Figure S3. Optogenetic silencing of MLIs does not disrupt CS suppression of SSs in neighboring cells, related to Figure 1.**

CKit cre mice were injected with 250 nl of AAV9-Ef1a-DIO eNpHR 3.0-EYFP nine sites in the cerebellum. Experiments were conducted 2-3 weeks later (see Methods). Slices were cut in order to evaluate expression and or ability to suppress MLI firing (A, B).

(A) Image of eNpHR 3.0-EYFP labelling showing an expression pattern that is characteristic of membrane labelling of MLIs, including the pinceaus associated with basket cells.

(B, C) On cell recordings were used to assess the effect of light on MLI firing, and we found that firing was eliminated in all MLIs.

After waiting for 2-3 weeks, *in vivo* recordings proceeded similarly to experiments shown in Figure 1. Once a pair was located, 5 s illumination was alternated with 5 s of no light, and this continued for at least 30 minutes in order to record sufficient complex spikes for each condition.

(D) Light increased PC firing of a pair of closely spaced (25 µm) cells.

(E) CS induced decreases in SS firing rate are shown for control condition (no light, left) and when MLI firing was suppressed with light (right).

(F) The effects of light on PC firing is shown for another pair of cells (50 µm).

(G) CS induced decreases in SS firing rate are shown for control condition (no light, left) and when MLI firing was suppressed with light (right).

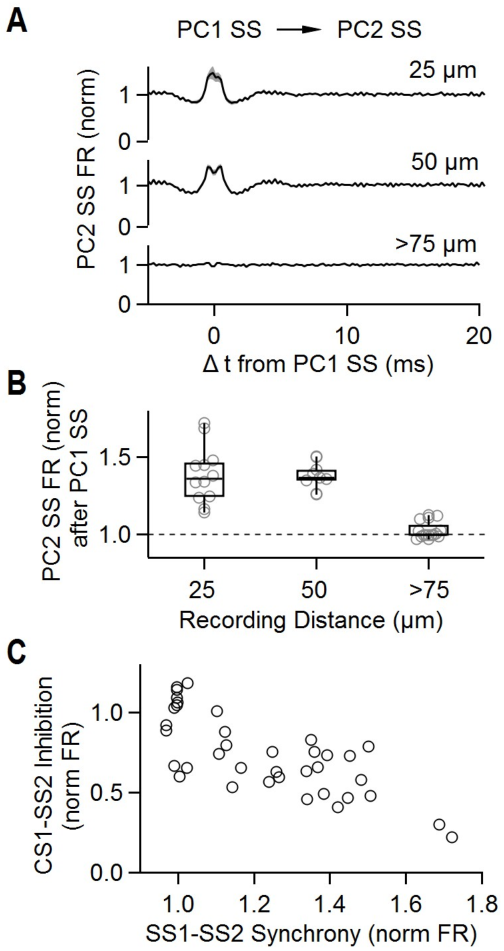


**Figure S4. Simple spikes promote synchrony whereas complex spikes suppress firing for neighboring cells, related to Figure 1.** The CC firing of the PC pairs of Figure 1 were analyzed as described previously (Han et al., 2018).

(A) Average firing rate of simple spikes from neighboring PCs (PC2 SS) after simple spikes from PC1 (PC1 SS). Recording sites were separated by 25 µm (top), 50 µm (middle), and more than 75 µm (bottom).

(B) Summary of normalized firing rates of PC2 SS after PC1 SS as a function of distance between recording sites.

(C) Summary of inhibition of PC2 SS by PC1 CS (from Figure 1) as a function of synchrony between PC1 SS and PC2 SS.

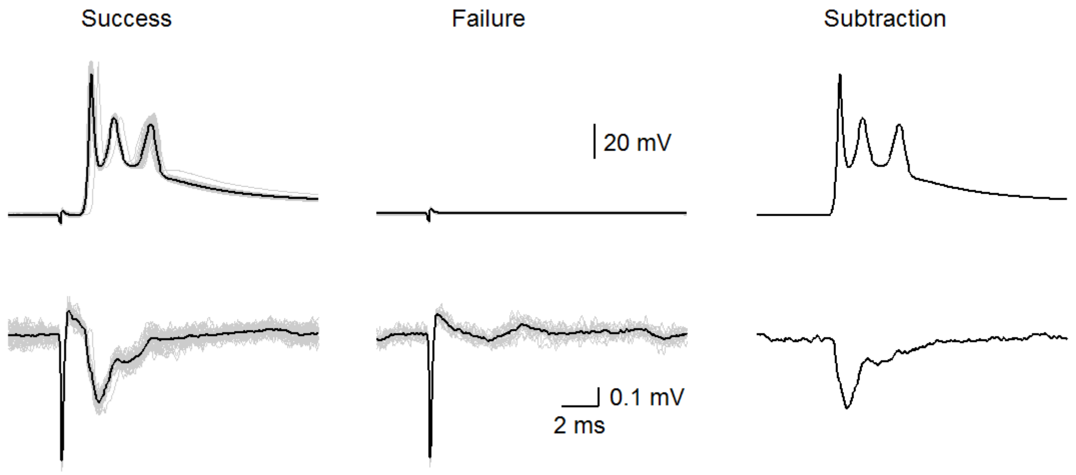


**Figure S5. Subtracted extracellular voltage responses by complex spikes, related to Figure 2.**
(top) Complex spikes by CF stimulation. A threshold stimulus intensity was used that stochastically evoked successes (left) or failures (middle) (individual trials: grey and average: black). Average success – average failure is shown (right)
(bottom) Extracellular signals near proximal dendrite by successful stimulation of the CF input (left), but no extracellular signals were observed when stimulation failed to evoke complex spikes (middle). Average success – average failure is shown (right).

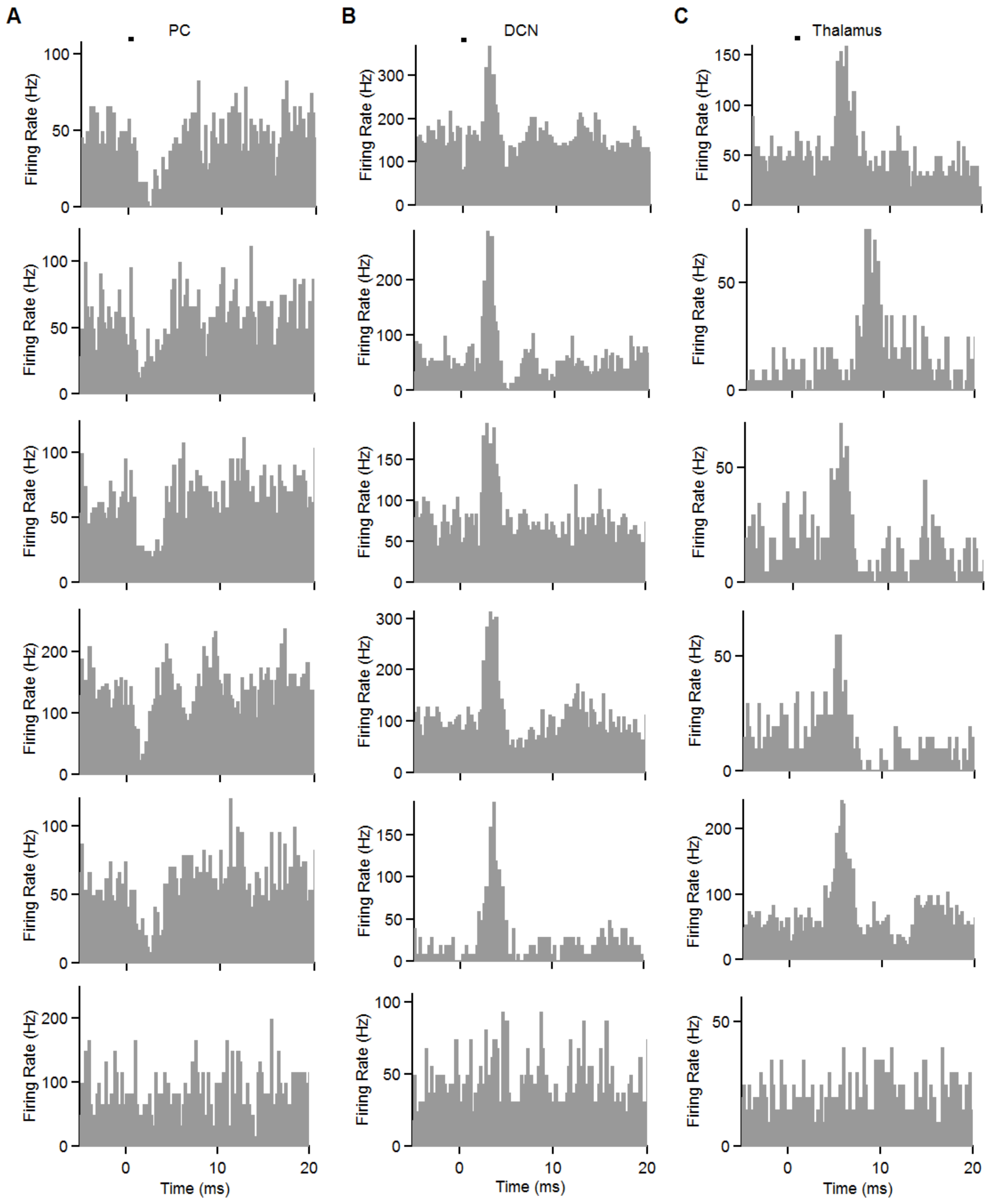


**Figure S6. Example light-evoked responses in PCs (A), DCN neurons (B) and in the motor thalamus (C), related to Figure 8.** The bottom row shows example cells in each area that did not respond significantly to stimulation.
